# Temporal adaptation aids object recognition in deep convolutional neural networks in suboptimal viewing scenario’s

**DOI:** 10.1101/2025.04.03.646752

**Authors:** Amber Marijn Brands, Georg Lange, Iris Isabelle Anna Groen

## Abstract

The primate visual system excels in recognizing objects under challenging viewing scenario’s. A neural mechanism that is thought to play a key role in this ability is rapid temporal adaptation, or the adjustment of neurons’ activity based on recent history. To understand how temporal adaptation may support object recognition, previous work has incorporated a variety of temporal feedback mechanisms in deep convolutional neural networks (DCNN) and explored how these mechanisms affect object recognition performance. While multiple adaptation mechanisms have been shown to impact model behavior, it remains unclear how the origin (intrinsic or recurrent) and the way the temporal feedback is integrated (additive or multiplicative) affects object recognition. Here, we compare the impact of four different temporal adaptation mechanisms on object recognition using three different task designs, including object recognition under either noise or occlusion, and in the context of novelty detection. Our results show that the effectiveness of temporal adaptation mechanisms for robust object recognition depends on the task and dataset. For objects embedded in noise, intrinsic adaptation excels with simple, high-contrast inputs, while recurrent mechanisms perform better with complex, low-contrast inputs, highlighting their focus on different visual features. Under dynamic occlusion, recurrent adaptation mechanisms exhibit a more progressive increase in performance over time, suggesting they better maintain object coherence when parts are obscured. For novelty detection, recurrent mechanisms show higher performance compared to intrinsic adaptation mechanisms, suggesting that recurrence aids in detecting global changes caused by the presentation of new objects. All together, these findings suggest that robust object recognition likely requires multiple temporal adaptation strategies in parallel to handle the diverse challenges of naturalistic visual settings.

## Introduction

The primate visual system can rapidly recognize objects even in challenging viewing conditions, for instance in noisy environments (Vinken et al., 2020; Brands et al., 2024), when objects are occluded (Zhu et al., 2019; Rajaei et al., 2019) and when novel objects are presented among familiar ones, referred to as novelty detection (Schomaker and Meeter, 2014; Jacob and Huber, 2020). In natural, real-world visual settings, objects are perceived within a dynamical stream of inputs, and hence in an inherently temporal context. Therefore, one strategy of the visual system that is thought to contribute to its robustness is the exploitation of temporal regularities in the visual inputs through a mechanism known as temporal adaptation, which refers to the suppression of a neuron’s activity based on its sensory history (Kohn, 2007; Wyatte et al., 2014; Weber and Fairhall, 2019; Wu et al., 2023). Temporal adaptation has been proposed to play a key role in modeling how the brain integrates neural information over temporal contexts, as it efficiently decorrelates inputs over time (Barlow and Foldiak 1989; Wang et al. 2004; for a review see Atick 1992) and also reduces the salience of recently seen stimuli in sensory processing (Kohn, 2007; Solomon and Kohn, 2014; Vogels, 2016; Whitmire and Stanley, 2016). Moreover, previous work has shown that adaptation of neural responses to recently perceived stimuli emerges in a remarkably similar way across sensory modalities and across species, implying that neural adaptation is governed by fundamental and conserved underlying mechanisms (Carandini and Heeger, 2012; Whitmire and Stanley, 2016).

Deep convolutional neural networks (DCNNs) have emerged as powerful tools to model biological vision (Yamins and DiCarlo, 2016; Kietzmann et al., 2017; Cichy and Kaiser, 2019). To understand how temporal adaptation may help achieve robust primate vision, researchers have attempted to endow feedforward DCNNs with computational mechanisms which take into account the temporal context and study the effect on object recognition (Tripathi et al., 2016; Beery et al., 2020; Spoerer et al., 2020). These computational mechanisms give rise to temporal adaptation either through intrinsic computations, operating on the level of the individual unit, or through recurrent computations. One recent study by Vinken et al. (2020) demonstrated that a DCNN implemented with an intrinsic adaptation mechanism can account for a wide range of neurophysiological and perceptual phenomena, and with its few trainable parameters, offers a less complex solution for biological systems than a recurrent network mechanism. They however showcased this for a DCNN trained on a single task optimized for one specific dataset, and it is therefore not clear whether these findings generalize to other tasks and datasets. Here, we test whether intrinsic adaptation mechanisms implemented in a DCNN are sufficient to achieve robust performance for object recognition across a range of suboptimal viewing conditions, or whether more complex, recurrent adaptation mechanisms are required. To achieve this, we systematically compare multiple temporal adaptation mechanisms implemented in a feedforward DCNN to solve object recognition in several challenging viewing conditions. Specifically, we optimize networks to perform classification in three scenario’s which have been shown to benefit from temporal adaptation, namely object recognition under noise (Vinken et al., 2020; Brands et al., 2024), under occlusion (Ban et al., 2013) and in the context of novelty detection (Dragoi et al., 2002; Kohn, 2007; Whitmire and Stanley, 2016; Weber and Fairhall, 2019).

We study four different types of adaptation mechanisms, which differ in the way they integrate a temporal feedback signal along two dimensions. The first dimension is the origin of the temporal feedback signal, namely only from the unit itself or also from units originating from other feature maps. Here, computations on the individual-unit level represent neural processes arising from intrinsic biophysical mechanisms operating within individual neurons (Whitmire and Stanley, 2016), while computations across feature maps within the same network layer represent neural processing in a neural circuit, whereby information is processed within a region, roughly corresponding to lateral recurrence (Felsen et al., 2002; Teich and Qian, 2003; del Mar Quiroga et al., 2016; Westrick et al., 2016). The second dimension is the type of interactions used to integrate previous inputs, namely through additive or multiplicative modulation, and is motivated by a previous study which showed that multiplicative interactions yield higher generalization performance against spatial shifts in the input compared to additive interactions (Brands et al., 2024). Both additive and multiplicative interactions have been observed in neural data, in the form of, for instance, linear summation (Hubel and Wiesel, 1962) and response gain (Carandini and Heeger, 2012), respectively. We trained a number of DCNN instances with one of the four different adaptation mechanisms on three different tasks, after which we compared network performances with a baseline model without temporal adaptation to better understand the benefit of integrating temporal context during object recognition. Moreover, to study the relationship with the content of the visual inputs, we trained on different datasets which differ in their feature complexity (MNIST, fashion MNIST and CIFAR10).

Our results demonstrate that the suitable mechanism for achieving robust object recognition strongly depends on both the task and the dataset a network is optimized for. First, we show that, when objects are embedded in noise, DCNNs with intrinsic adaptation mechanisms perform better for simple, high-contrast inputs, whereas DCNNs with recurrent mechanisms yield better performance for more complex, low-contrast inputs. This suggests that these adaptation mechanisms aid in processing different features of the incoming visual information when performing object recognition. Second, when objects are hidden by a sliding occluder, we show that enforcing adaptation through recurrent, as opposed to intrinsic, mechanisms leads to a smooth improvement in network performance over time. This suggests that recurrence aids networks to become more effective at maintaining a coherent perception of objects resulting in more robust recognition. Third, when multiple objects are presented simultaneously, we show that DCNNs with intrinsic adaptation mechanisms perform worse compared to DCNNs with recurrent mechanisms, implying that recurrence aids in better detection of global changes that are the result of the presentation of novel objects. Overall, our findings suggest that intrinsic adaptation mechanisms alone are not sufficient for robust object recognition. Instead, it is favorable to employ multiple temporal adaptation strategies in parallel when dealing with a wide variety of challenging views posed by naturalistic settings.

## Materials and Methods

### Stimuli

Here, we performed a set of experiments to study how different mechanisms giving rise to temporal adaptation implemented in a DCNN influences the network’s ability to recognize objects under three challenging viewing conditions. For the generation of the stimuli in each experiment, we used images that contained objects belonging to one of three datasets, namely MNIST (LeCun et al., 2010), fashion MNIST (fMNIST, Xiao et al. 2017) or CIFAR10 (Krizhevsky et al., 2009) and are from here on out referred to as *target* images. To study the role of temporal adaptation during object recognition under suboptimal viewing conditions, we presented DCNNs with sample sequences, which contained a degraded form of the target image using either noise (**Fig. 1A**, *Experiment 1*) or occlusion (**Fig. 1A**, *Experiment 2*). Another set of experiments investigated object recognition in the context of novelty detection, whereby novel target images were added throughout the sequence (**Fig. 1A**, *Experiment 3*). Each experiment is described in more detail below.

**Figure 1:**
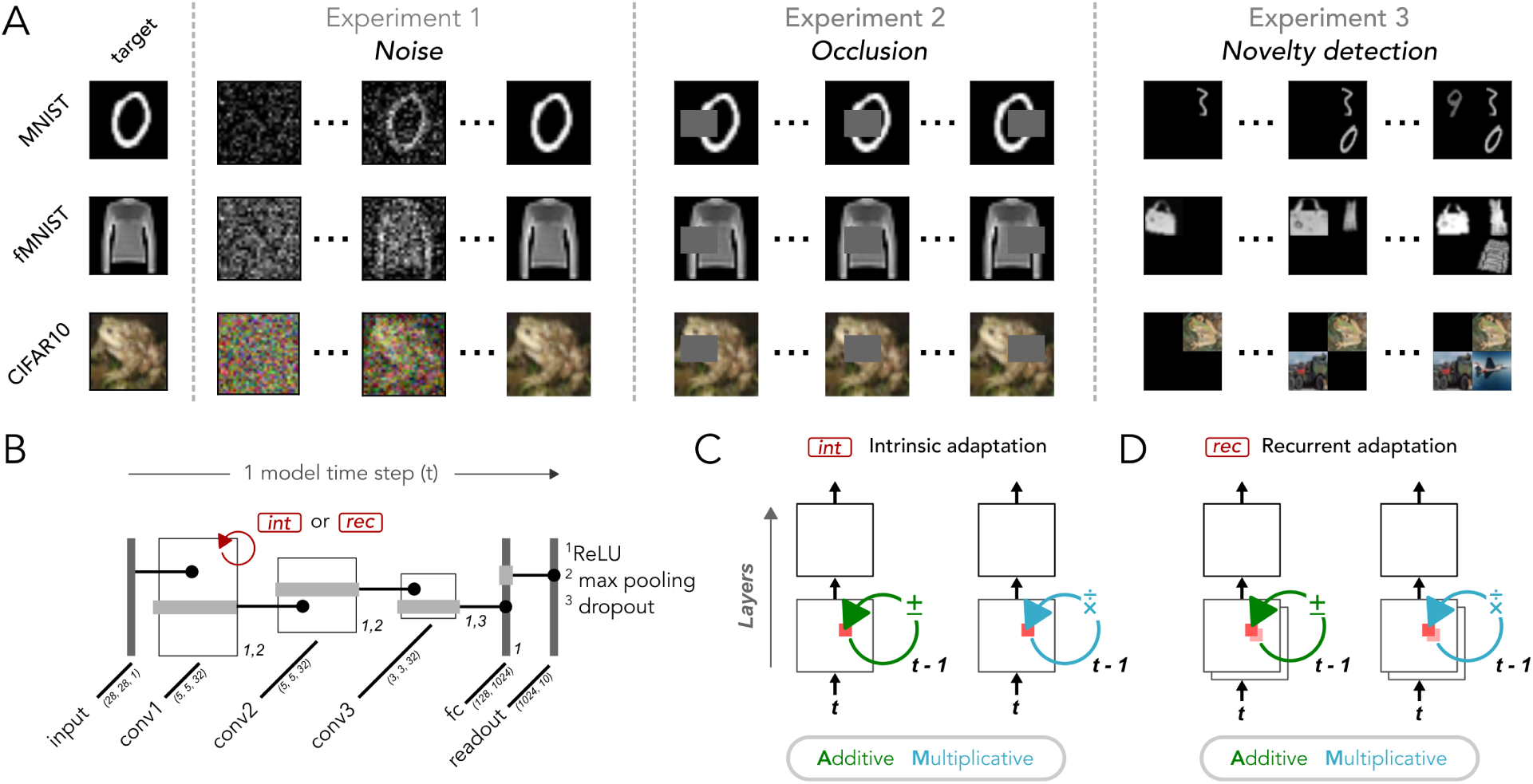
Experimental design and network modeling. **A**: Exemplar stimuli of the three tasks the DCNNs were trained on. Each row depicts an example sequence belonging to either the MNIST (*top*), fashion MNIST (*middle*) and CIFAR10 (*bottom*) dataset. *Experiment 1 Noise*, Object recognition with targets embedded in a temporally repeatedly noise pattern, with two types of noise manipulations (see **Supp. Fig. 1**). *Experiment 2 Occlusion*, Object recognition during which the target was occluded by a patch sliding over the image. *Experiment 3 Novelty detection*, Object recognition whereby the model was trained to categorize novel as opposed to familiar targets. **B**: Deep convolutional neural networks (DCNNs) were trained with a feedforward backbone consisting of three convolutional filter layers, one fully connected layer and a readout layer. After the first convolutional layer, temporal adaptation was applied (depicted in red) by integrating a temporal feedback signal using intrinsic or recurrent computations. The temporal feedback signal consisted of activations from the previous model time step, i.e. previous feedforward pass. Filter sizes are represented within parentheses. **C**: Intrinsic adaptation mechanisms which operate on the level of the individual unit. Left, An additive form introduced previously by Vinken et al. (2020), whereby unit activations from the previous timestep are added to or subtracted from the same unit’s activation at the current timestep. Right, A multiplicative form introduced by Heeger (1992, 1993) whereby activations from the unit’s current timestep are multiplied or divided by the same unit’s activation at the previous timestep. **D**: Temporal adaptation mechanism which feed unit activations across feature maps either through additive (left) or multiplicative (right) interactions, inspired by Vinken et al. (2020).

### Experiment 1: Object recognition under noise

#### Experiment 1.1

First, we aimed to replicate and confirm the results of Vinken et al. (2020), showing that intrinsic adaptation mechanisms operating on the individual unit level are sufficient to obtain robust object recognition under noise. To achieve this, we created similar image sequences as were used in their study, whereby target images were embedded in a temporally repeated noise pattern. Inputs samples consisted of three sequential images equivalent to three timesteps (for example sequences see **Supp. Fig. 1A**). The first image, referred to as the *adapter*, contained pixels drawn from a Gaussian distribution (*µ* = 0, *σ* = 0.2) at the same size and resolution as the input image. The second image in the sequence was blank, whereby all pixel values were set to the mean of target image. This was to ensure consistency in the mean luminance of the sequence and to prevent abrupt transitions in brightness. The third image, referred to as the *test* image, combined the same noise pattern as used in the adapter image with the target image, which was to be classified. To vary task difficulty, we randomly altered the contrast level for each individual sequence to either 20% or 100%, whereby the contrast adjustment was performed as follows:

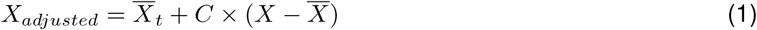

where *X_adjusted_* represents the contrast-modified target image, *X* the mean pixel value of the original target image, *C* the desired contrast level (i.e. 0.2 and 1.0 for a contrast of 20% and 100%, respectively) and *X* is the original target image. The adapter and test image were clamped to the range of [0, 1].

We tested the networks on unseen data using five different test sets. These test sets were designed to evaluate the networks’ robustness to variations in the visual input that could arise in real-world scenarios (e.g. spatial variation, noise distribution) and to understand how each adaptation mechanism might facilitate recognition of objects. Three of these were introduced earlier by Vinken et al. (2020): two contain the same Gaussian noise as the training set but with either different variances or means (range of 0 and 1 with a step size of 0.2) and one contains uniform instead of Gaussian noise with different variances (range of 0 and 1 with a step size of 0.2). The fourth test set had the same noise as used during training, but for the test image, the noise pattern was spatially shifted to the right with a varying number of pixels (ranging from 1 to 7). Lastly, the fifth test set had again the same type of noise as used during training, but the spatial distribution of pixel values was different during the adapter and test image.

#### Experiment 1.2

To study object recognition during noise in a more challenging viewing setting, we created a new stimulus set, whereby instead of abruptly adding the noise, as was the case for Experiment 1.1, the contribution of the noise was gradually decreased, thereby making the target image gradually more visible with more model timesteps. This results in a trade-off between suppression of the noise and keeping units responsive to the emerging target image.

Sequences consisted of 10 images (for exemplar sequences see **Supp. Fig. 1B**). Here, the first image consisted, same as Experiment 1.1., of a noise pattern sampled from a Gaussian distribution. Subsequently, in the following timesteps, the noise was combined with the target image according to the following equation:

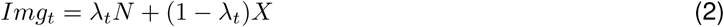

where *Img_t_* represents the image at timestep *t*, *N* represents the noise pattern, *X* represents the target image and *λ_t_* represents a weight factor at time *t*, which was set to 0.9 at *t* = 0 and decreased with a fixed value of 0.1 after each timestep. The last image of the sequence had a weight factor of 1 and depicted the target image without noise. The values were clamped to the range of [0, 1] at each timestep.

### Experiment 2: Object recognition under occlusion

Another suboptimal viewing condition in the context of object recognition is when objects are occluded. In naturalistic settings, occluders often move across an image, whereby different parts of the object becomes visible or hidden in a progressive manner.

To mimic these spatio-temporal dynamics in naturalistic visual inputs, we created sequences with an occluder, consisting of a gray patch, placed on the left or right side of a target image which shifted horizontally with every model timestep. The number of images in a sequence varied by dataset, with MNIST or fMNIST consisting of 9 and CIFAR10 of 11 images within one sequence (for exemplar sequences see **Supp. Fig. 2**). This difference was due to CIFAR10’s larger image size (32 pixels vs. 28 for MNIST and fMNIST) and the fact that the patch covered the entire image width with a fixed number of pixels. The size of the occluder was set to 12 × 12 pixels, which ensured that some part of the object was hidden, but enough of the object was still visible to leverage spatiotemporal cues in the object features (though initial exploration indicated results were robust over different pixel sizes). To introduce some variability in the spatial location of the occluder, we randomly varied the vertical starting position of the patch. At each timestep inputs were clamped between a value range of [0, 1].

To understand how temporal adaptation aids in the recognition of the target when hidden by a sliding occluder, we let the networks perform classification for every image in the sequence, yielding recognition performance at each model timestep. Ultimately, this informs us how the different temporal adaptation mechanisms handle the dynamical occlusion and to what degree they maintain a coherent representation of the object over time when different parts are hidden.

### Experiment 3: Novelty detection

We also examined the role of temporal adaptation in the context of novelty detection, whereby human studies have shown that salience is higher for novel compared to familiar inputs (Sokolov, 1963; Corbetta and Shulman, 2002). For this, we created image sequences of 20 model timesteps whereby novel target images were dynamically added to one of four quadrants (for exemplar sequences see **Supp. Fig. 3**). The size of each quadrant was set to the size of the target image of the respective dataset, resulting in a total image size of 56 × 56 pixels for the MNIST and fMNIST and 64 × 64 pixels for the CIFAR10 dataset. Every sequence was structured to include at least one target that appeared in the very first image. After the first timestep, appearance (or “onset”) of the other target images was randomized in terms of the position (i.e. quadrant) and timestep of the onset. After onset, the location of a given target remained constant throughout the input sequence. Moreover, to determine to what degree the different temporal adaptation mechanisms can distinguish local changes to familiar objects from global changes related to the presentation of novel objects, we applied a set of image augmentation techniques (e.g. rotation, scaling) resulting in small variations in object appearance over model timesteps. For the details regarding the different augmentation techniques see **Supplementary Table 1**. Lastly, at each timestep inputs were clamped between a value range of [0, 1].

To study the role of different temporal adaptation mechanisms during novelty detection, networks were trained to predict the object belonging to the most recently introduced target image. In cases where no target was added at a timestep, the model’s objective was to identify the most recently added target. If multiple targets appeared simultaneously in a timestep, the model was trained to predict the target with the highest contrast. Overall, this task design allowed us assess to what degree different temporal adaptation mechanisms aid in adapting to changes of familiar targets undergoing local variations and detecting global changes as a result the presentations of novel inputs.

### Computational modelling

#### Models

Deep neural networks were implemented using Pytorch (Paszke et al., 2019) with standard convolutions and linear layers. For models trained on Experiment 1 (noise) and Experiment 2 (occlusion), three convolutional layers were followed by one fully connected and one readout layer (**Fig. 1B**), whereby the fully connected layer had a dimensionality of 1024, which was reduced by the readout layer to 10 output units. For MNIST and fashion MNIST trained models, the convolutions had 32 feature maps each and a kernel size of 5, 5, 3 for the first, second and third layer, respectively. For CIFAR10 models, inputs were more complex (three input channels instead of one). To boosts the network’s representational power we increased the number of feature maps to 32, 64 and 64. For Experiment 3 (novelty detection), inputs were also more complex (larger input size) and for this reason we added a fourth convolutional layer between the last convolution and the fully connected layer with 32 (MNIST and fashion MNIST) or 64 (CIFAR10) feature maps and a kernel size of 3. For a detailed overview of model architectures see **Supplementary Table 2**.

We introduced temporal adaptation into the networks by feeding activity from previous timesteps, with one timestep *t*, defined as one feedforward sweep. We define a linear response **L***_n_*(*t*) at time *t* as the output of a convolutional layer:

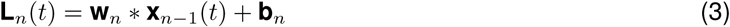

given the unit’s current input **x***_n−_*_1_(*t*), bottom-up convolutional weights **W***_n_* and biases **b***_n_*, whereby the convolution operation is represented as ∗. Five different model types were trained, with five model instances per task and dataset. For the baseline model, no adaptation mechanism was added such that all images were processed independently over model time steps. For the remaining models, we implemented and compared four different temporal adaptation mechanisms (see below). Deviating from Vinken et al. (2020) and Brands et al. (2023), we only apply adaptation in a single layer, which allows us to better understand and interpret the emerging dynamics, essentially mimicking adaptation in early visual areas only. Corresponding temporal adaptation parameters were optimized simultaneously with the feedforward DCNN parameters. An overview of the number of parameters for each network architecture and dataset is provided in **Supplementary Table 4** and the temporal adaptation parameter values obtained after training are reported in **Supplementary Figure 4**-7.

#### Intrinsic temporal adaptation

*Additive intrinsic adaptation.* We implemented an additive from of intrinsic adaptation that emerges from the properties of individual units (**Fig. 1C**, *left*), as introduced previously by Vinken et al. (2020). Here, each unit *i* in the network has an exponentially decaying intrinsic adaptation state, *s_t_*, which is updated at each time step *t*, based on its previous state *s*(*t* − 1) and the previous response *r*(*t* − 1) as follows:

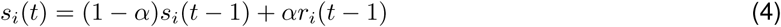

where *α* is a constant determining the time scale of the decay. This intrinsic adaptation state is then subtracted from the linear response before applying the activation function *ϕ*.

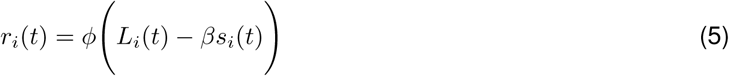

where *β* is a constant scaling the amount of suppression. For *β >* 0, these updating rules result in an exponentially decaying response for constant input that recovers in case of no input.

*Multiplicative intrinsic adaptation.* For the multiplicative form of intrinsic adaptation, we take inspiration from previous work (Brands et al., 2024), which endowed networks with divisive normalization. Divisive normalization was originally proposed by Heeger (1992) and has been studied extensively in the spatial domain. Here, we apply this to the temporal domain (**Fig. 1C**, *right*) by implementing a mathematical framework originally formulated in Heeger (1992, 1993), which defines a neuron’s response *r_i_* recursively over time. For each unit *i* in the network the response at timepoint *t* is updated before applying the rectifier activation function *ϕ* such that:

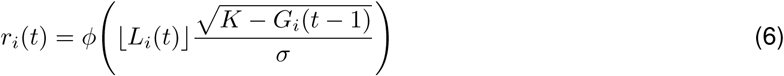

where *G*(*t* − 1) is a temporal feedback signal from the previous time step which is updated based on its previous state and the current response:

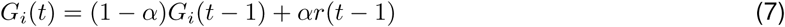

where *α* determines the time scale of the feedback signal. This multiplicative feedback signal results in divisive suppression (for details, see Heeger 1993, Appendix A).

#### Recurrent temporal adaptation

In addition to temporal adaptation arising from intrinsic biophysical mechanisms inside a single neuron, temporal adaptation phenomena have also been proposed to be the result of recurrent interactions between neurons. Here, we incorporated such a form of temporal adaptation using a method adapted from Vinken et al. (2020), inspired by computational models which implemented adaptation by recurrent interactions between orientation tuned channels (Felsen et al. 2002; Teich and Qian 2003; Westrick et al. 2016; **Fig. 1D**, *left*), as follows:

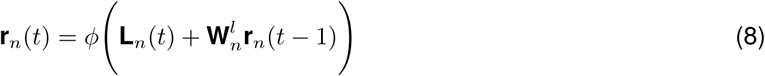

where **W***^l^* are the lateral weights consisting of 32 kernels of size 1 × 1 × 32 (stride = 1). To compare additive and multiplicative interactions, we also implement a multiplicative form of recurrent adaptation (**Fig. 1D**, *right*):

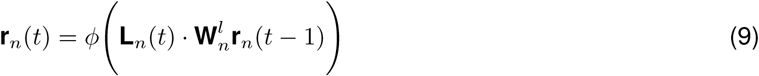

with the same lateral weights (i.e. 1 × 1 × 32) as used for the additive form.

To summarize, we explore these four adaptation mechanisms, which differ in two key dimensions: the origin of the temporal feedback (intrinsic or recurrent) and the type of interaction used to integrate previous inputs (additive or multiplicative). Note that even though the multiplicative form of recurrent temporal adaptation does not directly correspond to the multiplicative form of intrinsic adaptation, i.e. divisive normalization, the motivation for the formulation for this recurrent form was to make the mechanism mathematically consistent with the intrinsic mechanism in terms of how the feedback is integrated.

#### Training procedures

For Experiment 1 (noise) and Experiment 2 (occlusion), models were trained for 20 epochs, whereby a class prediction was made at the last timestep for Experiment 1.1 and at all timesteps for Experiment 1.1 and Experiment 2. For Experiment 3 (novelty detection) models were trained for 10 epochs, whereby a class prediction was made at every timestep. Each epoch contained 60.000, 60.000 and 50.000 sample sequences for the MNIST, fashion MNIST and CIFAR10 dataset, respectively, whereby targets were sampled from the training set. For the optimization we made use of the Adam optimization algorithm, a learning rate of 0.001, a batch size of 128 and 50% drop-out after the last convolutional layer. All models converged well before training ended, with the exception of the models used for Experiment 3 trained on the CIFAR10 dataset, which were therefore omitted from analysis. For a detailed overview of the training procedures see **Supplementary Table 3**. The results presented in **Figure 2-5** include analyses using image sequences with target images sampled over the datasets’ test set (10.000 for all datasets).

**Figure 2:**
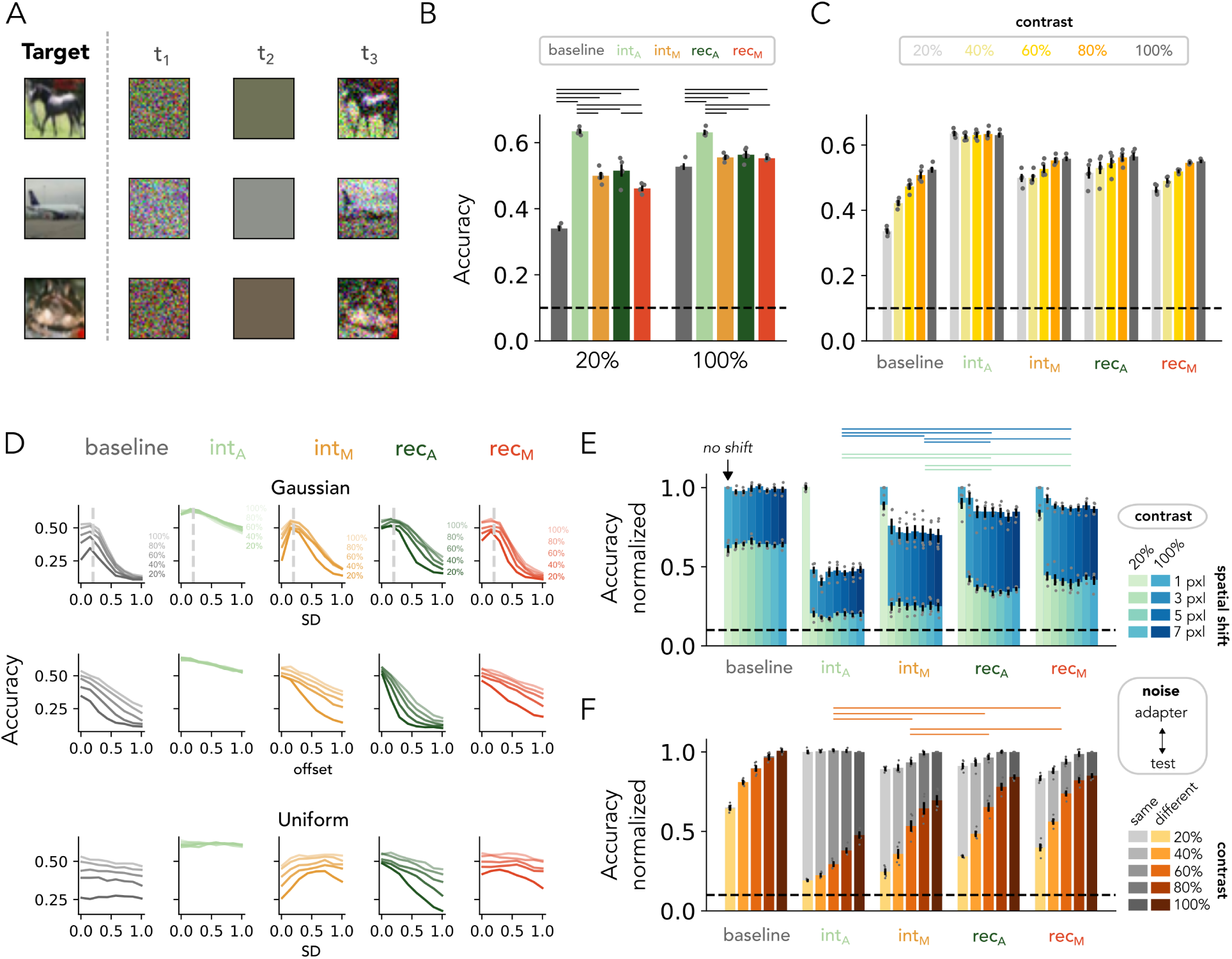
DCNNs with additive intrinsic adaptation solves task by subtracting the noise. **A**: Each row depicts an example sequence. The first image in the sequence (*t*_1_) contains Gaussian noise and is followed by a blank image (*t*_2_). The third image (*t*_3_) contains a CIFAR10 target image embedded in the noise pattern presented at *t*_1_. **B**: Test accuracy for models with and without (*baseline*) temporal adaptation trained on a mix of high- (100 %) and low- (20%) object contrasts. DCNNs with additive intrinsic adaptation (*int_A_*) exhibit highest performance for both object contrast levels. **C**: Model performance when generalizing to different object contrasts (only the contrast levels in gray were included in the training set). Superior performance of DCNNs with additive intrinsic adaptation (*int_A_*) consistent across object contrast levels. **D**: Model performance when generalizing to different noise patterns (see Materials and Methods, *Experiment 1: Object recognition under noise*). Superior performance of DCNNs with additive intrinsic adaptation (*int_A_*) consistent across different types of noise. **E**: Model performance when spatially shifting the noise pattern during test. The first bar for each adaptation mechanism depicts performance without a spatial shift. DCNNs with additive intrinsic adaptation (*int_A_*) show poorest robustness against spatial shifts. **F**: Model performance when the test image (*t*_3_) contains a different noise pattern than the adapter image (*t*_1_). Grey bars mark accuracy when tested on the same noise pattern (same data as shown in panel C). DCNNs with additive intrinsic adaptation (*int_A_*) show poorest robustness against modifying the noise during test. For panel B, C, E and F, the dashed line depicts chance level (i.e. 10%), markers represent individual network initializations and the error bars depict the SEM across these network initializations. For panel B, E and F, horizontal stripes on the top of the plot indicate significant differences between networks (One-way ANOVA, post-hoc Tukey test, *p <* 0.05).

### Data and code availability

All code used for the purpose of this paper can be found at the GitHub repository https://github.com/ ABra1993/tAdaptation_DNN.git.

## Results

In this study, we investigate how biologically-inspired temporal adaptation mechanisms contribute to robust object recognition in suboptimal viewing scenarios. To achieve this, we enhanced a feedforward DCNN with temporal adaptation capabilities by incorporating a temporal feedback signal through one of four computational mechanisms (see Materials and Methods, *Models*; **Fig. 1B-D**). Two of these mechanisms, namely additive intrinsic adaptation (see *Eq. 5*) and additive recurrent adaptation (see *Eq. 8*), comprise a feedback signal which originates from the unit itself or from other feature maps as well, respectively, and is incorporated through additive interactions. The other two mechanisms use multiplicative rather than additive modulation to integrate the temporal feedback signal, again either through intrinsic (see *Eq. 6*) or recurrent adaptation (see *Eq. 9*). Below, we examine how each temporal adaptation mechanism contributes to robust object recognition by optimizing the DCNNs on three different tasks that included manipulations associated with real-world scenario’s, including object degradation under noise and occlusion as well as in the context of novelty detection during multiple-object presentations.

### Additive intrinsic adaptation leads to robust object recognition under noise

A challenging scenario for object recognition which may benefit from temporal adaptation is during noise. For example, during the recognition of a car on a misty road. We first replicate and confirm previous findings by Vinken et al. (2020), who showed that a simple, intrinsic adaptation mechanism at the individual unit level is sufficient for achieving robust object recognition in the presence of noise. DCNNs were trained on an object classification task whereby objects were embedded within a temporally repeated noise pattern (**Fig. 2A**). We expand the approach by Vinken et al. (2020) and in addition to high contrast objects, present objects with low contrast to assess how temporal adaptation mechanisms perform under varying levels of noise degradation. Here, temporal adaptation aids in robust object recognition by allowing the DCNN to adapt to the noise pattern, leading to suppression of the noise over time and thereby enhancing the visibility of the object. We here report results on CIFAR10, as these showed the most divergent performance for the different temporal adaptation mechanisms. Results for the other two datasets, MIST and fashion MNIST, can be found in **Supplementary Figure 8** and **Supplementary Figure 9**, respectively.

The baseline model without temporal adaptation achieves an accuracy of around 50% for the high-contrast images (with a chance level of 10%), which drops considerably to around 35% for low-contrast images (**Fig. 2B** and **Supp. Table 5**), illustrating the challenge of the task to separate a low-contrast signal from a noisy background. In contrast, DCNNs endowed with temporal adaptation show higher performance than the baseline model, with interestingly more gain in accuracy for low than high contrast images (20% contrast, one-way ANOVA, *F*_4_ = 124.56, *p <* 0.001; 100% contrast, one-way ANOVA, *F*_4_ = 39.24, *p <* 0.001). This indicates that the models utilize the adaptation mechanism to suppress the task-irrelevant inputs (i.e. the noise), thereby considerably raising the signal-to-noise ratio for the target images. Moreover, in line with the findings of Vinken et al. (2020), the DCNN with additive intrinsic adaptation (*int_A_*) exhibits highest performance and least deterioration as a result of lowering the contrast of the object image. Performance of networks with additive intrinsic adaptation also generalize well to a broader range of contrasts (**Fig. 2C**), while DCNNs with multiplicative intrinsic and recurrent adaptation show a gradual decrease when the object contrast is lowered. To further assess model ability to generalize to unseen inputs, we adapted the approach of Vinken et al. (2020) and we tested how well the models perform when the distributions of noise patterns is different during training and testing. We find that DCNNs with additive intrinsic adaptation also generalize best to higher standard deviations of Gaussian noise (**Fig. 2D**, *top*), Gaussian noise with an offset (**Fig. 2D**, *middle*), and uniformly distributed noise (**Fig. 2D**, *bottom*). These results are consistent across datasets, namely MNIST (**Supp. Fig. 8B-D** and **Supp. Table 6**) and fashion MNIST (**Supp. Fig. 9B-D** and **Supp. Table 7**). Together, these findings show that a simple additive intrinsic adaptation mechanism achieves robust object recognition and exhibits greater resilience to variations in contrast and noise compared to other temporal adaptation mechanisms, including multiplicative intrinsic and recurrent adaptation.

### Additive intrinsic adaptation solves object recognition by subtracting the noise

So far, we have confirmed prior findings that a DCNN with a simple, additive intrinsic adaptation mechanism can achieve robust object recognition in temporally repeated noise patterns. However, one possible reason for the superior performance of the intrinsic adaptation mechanism is that the task design, as introduced by Vinken et al. (2020), may be particularly suited for a solution where the DCNN learns to simply subtract the noise from the test image. Given that each training sample consists of only three images (**Fig. 2A**), the adaptation state during the test image is strongly influenced by the adapter image, making it easier for the network to perform a straightforward subtraction at the individual unit level.

To investigate if the superior performance of DCNNs with additive intrinsic adaptation indeed arises from simple subtraction of the noise, we presented the networks with a test set which assesses the networks’ generalizability to other input conditions: Instead of changing the noise patterns we test the network on, we use the same noise patterns the model was trained on but spatially shift it at the test stage (t_3_). The test image now still has the same noise pattern but with a slight offset such that unit-wise subtraction is compromised. Results show that even when shifting the noise pattern only by a single pixel, DCNNs with additive intrinsic adaptation (*int_A_*) exhibit a drop in performance of about 54% for high object contrasts and an even larger drop of 81% for low object contrasts, suggesting that the model relies on complete subtraction (**Fig. 2E**). In contrast, the other adaptation mechanisms show significantly higher robustness against spatial shifts in the noise (low contrast, one-way ANOVA, *F*_4_ = 124.56, *p <* 0.001; high contrast, one-way ANOVA, *F*_4_ = 39.24, *p <* 0.001).

We also evaluated the behavior of the models when we resample the noise pattern at the test stage, such that adapter and test noise differ. This again revealed a more pronounced reduction in accuracy for DCNNs with additive intrinsic adaptation compared to the other adaptation mechanisms (averaged over all contrasts, one-way ANOVA, *F*_4_ = 168.30, *p <* 0.001; **Fig. 2F**). These results are overall consistent across datasets (MNIST, **Supp. Fig. 8D-F**; fashion MNIST, **Supp. Fig. 9D-F**), though we observe that MNIST-trained DCNNs with additive recurrent adaptation also show poorer robustness, while DCNNs with multiplicative adaptation mechanisms are less affected by the spatial shifts (**Supp. Fig. 9F**). Note that the baseline model is not affected by the noise modification, which can be explained by the fact that it does not adapt to previous inputs and processes each image independently. Overall, these findings reveal that DCNNs integrating temporal context using additive interactions simply learn to subtract the noise, implying that the benefit of this additive modulation on robust object recognition depends on the specific task design.

### Benefit of temporal adaptation depends on input complexity

In the previous section, we have revealed a limitation of the task design by Vinken et al. (2020): there is no incentive for the models to regulate the strength of suppression, unlike biological systems where adaptation is finely tuned in both strength and duration to optimize visual perception over time. For instance, recognizing a car on a misty road requires adapting to the irrelevant noise from the mist, while remaining responsive to changes in input, such as the approaching car gradually becoming visible. We expect that simple subtraction will not be sufficient: given that the object is present across multiple time steps, suppression could result in the units becoming overall less responsive to novel inputs, which in this case belong to the object. This creates a trade-off between suppression of features evoked by the noise on the one hand, and keeping units responsive to serve recognition on the other. To better mimic these real-world scenario’s, we modified the previous task, whereby instead of adding the noise pattern solely at the last timestep, we gradually let it decrease by re-weighting the contribution of the target image and noise pattern with every model timestep (**Fig. 3A**; see Materials and Methods, *Experiment 1: Object recognition under noise*).

**Figure 3:**
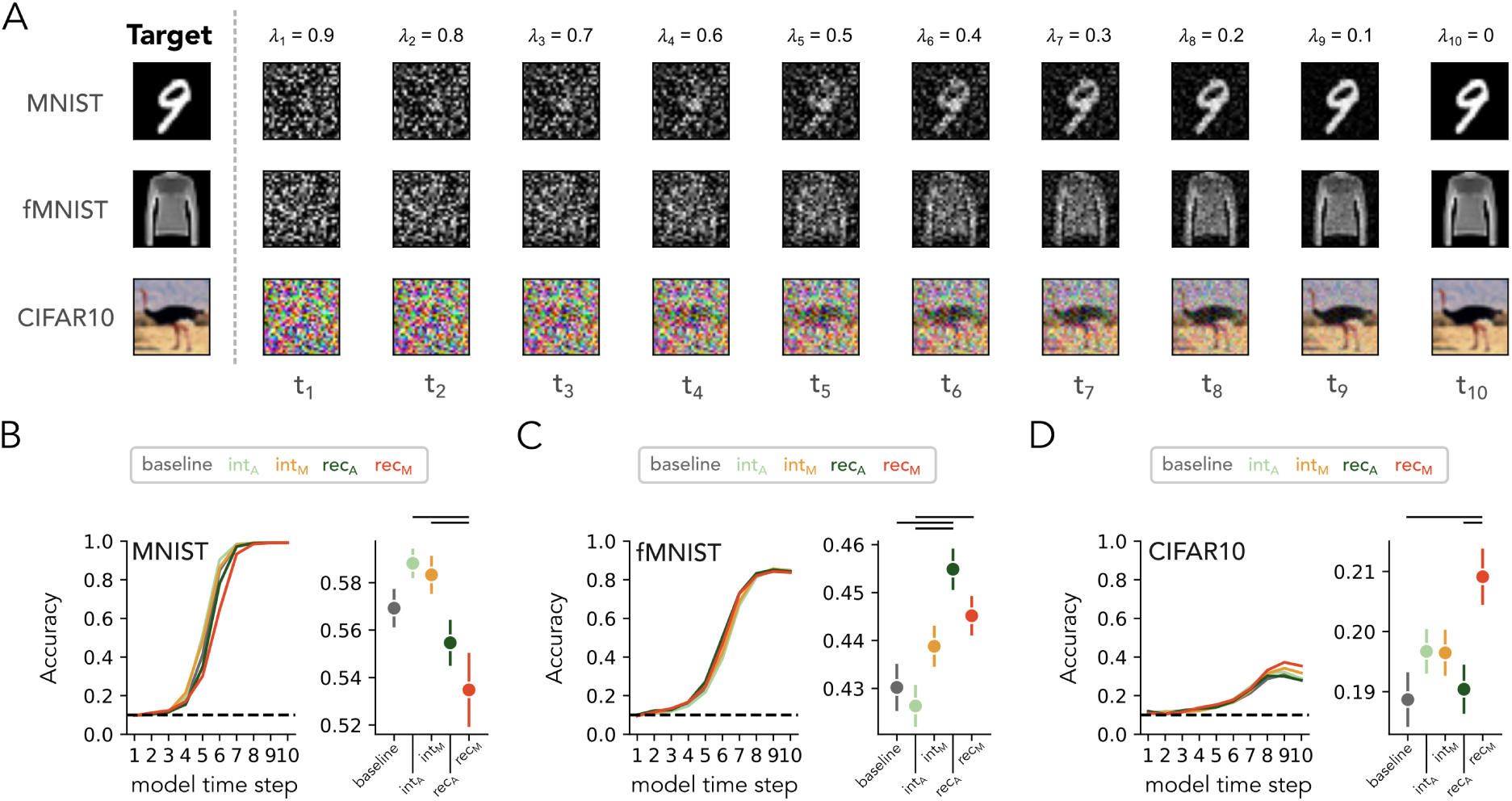
Benefit of temporal adaptation during object recognition under gradually decreasing noise is dataset-dependent. **A**: Each row depicts an example sequence consisting of an object-containing target image interleaved with noise, whereby the contribution of the noise, *λt*, decreases a fixed amount with every timestep. **B**: *Left*, Classification accuracy for models without (baseline) and with a temporal adaptation mechanism. *Right*, Classification accuracy averaged over model timesteps. Values depict means, error bars depict SEM across network initializations and the horizontal gray lines depict significant differences between networks (one-way ANOVA, post-hoc Tukey test, *p <* 0.05). **C-D**: Same as (B) for networks trained on the fashion MNIST (C) and CIFAR10 (D) dataset.

In this more challenging task design, there is no longer a consistent advantage of intrinsic adaptation on DCNN performance. Instead, the best-performing temporal adaptation mechanism depends on the dataset the networks were optimized for (**Fig. 3B-D** and **Supp. Table 5-7**). MNIST-trained DCNNs with intrinsic adaptation, both the multiplicative and additive form, show higher performance compared to recurrent mechanisms, especially the multiplicative form (one-way ANOVA, *F*_4_ = 4.74, *p* = 0.003; **Fig. 3B**), seemingly outperforming the baseline model, though the pairwise difference is not significant (int*_A_*, *F*_4_ = 0.019, *p* = 0.67; int*_M_*, *F*_4_ = 0.014, *p* = 0.86). Fashion MNIST-trained DCNNs with recurrent adaptation, and in particular the additive form, outperform both the baseline model and the models with intrinsic adaptation (one-way ANOVA, *F*_4_ = 7.01, *p <* 0.001; **Fig. 3C**). For the CIFAR10-trained DCNNs, we observe overall poorer model performance (**Fig. 3D**), whereby DCNNs with multiplicative recurrent adaptation outperform the other models only at the last few timesteps (one-way ANOVA, *F*_4_ = 3.69, *p* = 0.01). These results show that intrinsic adaptation mechanisms perform better for datasets with low complexity (i.e. MNIST) and recurrent mechanisms better perform for more complex datasets (fashion MNIST and CIFAR10)

The divergence of results across datasets suggests that the potential benefit of a given adaptation mechanism may depend on specific features in the visual input: intrinsic adaptation mechanisms, operating at the individual unit level without interaction across feature maps, perform better on simpler, high-contrast datasets with strong edge-like features, like MNIST. In contrast, recurrent mechanisms, which integrate information across feature maps, perform better on more complex, low-contrast datasets with varying textures, such as fashion MNIST and CIFAR10. Overall, optimization on a task design that does not solely favor response suppression shows that intrinsic mechanisms excel at processing simple, high-contrast features, while recurrent mechanisms are better equipped to handle complex, low-contrast features, ultimately enhancing the networks’ ability to recognize objects under noise.

### Recurrent interactions contribute to robust recognition during dynamic occlusion

Another challenging scenario for object recognition which may benefit from temporal adaptation is during dynamic occlusion. In naturalistic settings, when an occluder moves in front of a scene, parts of the object become visible or hidden in sequence. For example, when walking past a tree, different portions of a car parked behind it are gradually obscured and revealed as the view changes. In this scenario, temporal adaptation is useful as it allows for the adjustment to the shifting visibility of the object over time and integrate information from partially revealed parts of the object (Berzhanskaya et al., 2007; Battaglini et al., 2015). To examine the impact of the different temporal adaptation mechanisms on object recognition under such spatio-temporal dynamics, we conducted an experiment whereby the object in the target image is occluded by a rectangular patch which horizontally slides across the target image across model timesteps (**Fig. 4A**; see Materials and Methods, *Experiment 2: Object recognition under occlusion*). This sliding patch paradigm requires the network to update its representations dynamically as new portions of the image become available.

**Figure 4:**
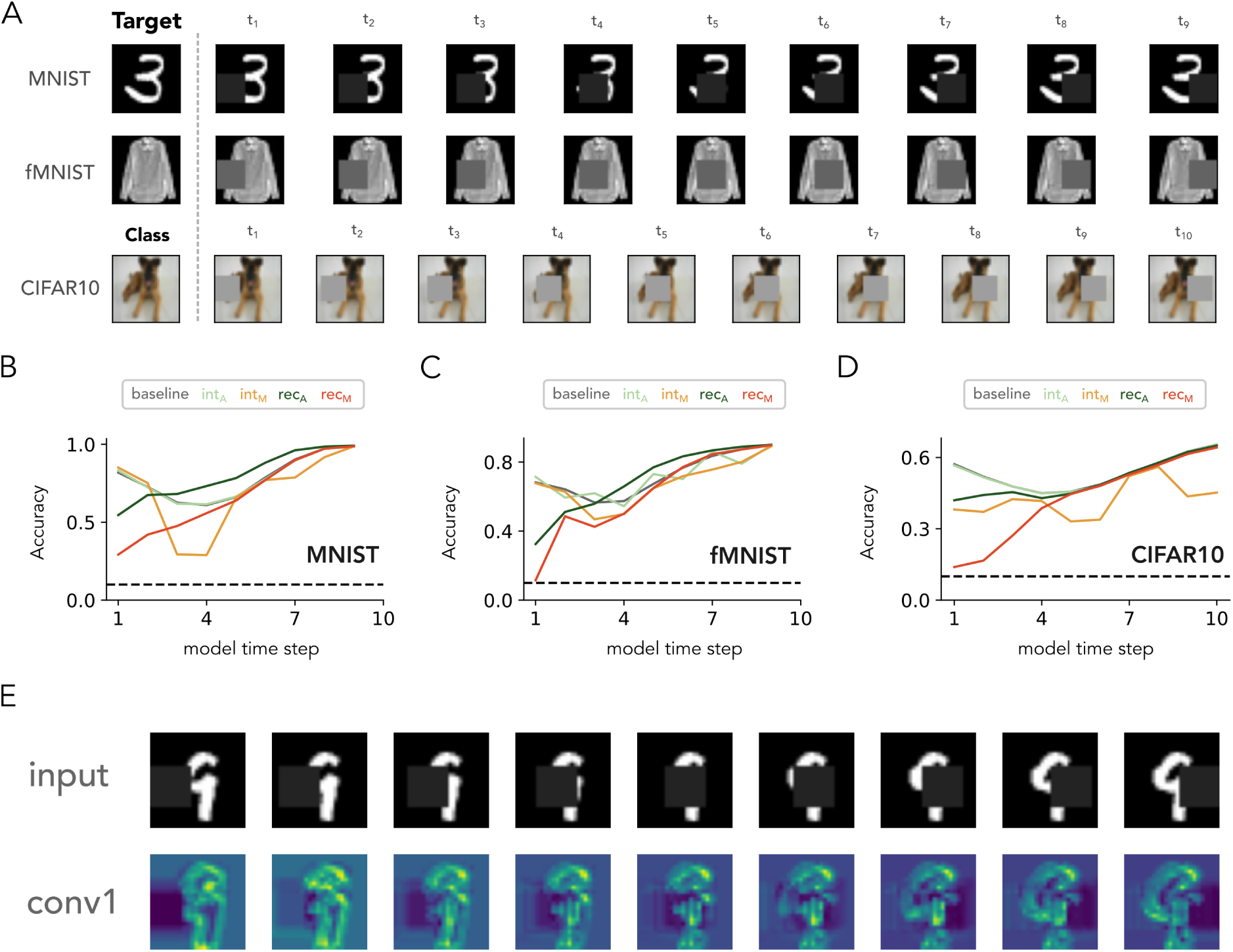
Recurrent adaptation mechanisms result in a smooth and consistent improvement in performance over time. **A**: Input sequences consisting of objects belonging either to the MNIST, fashion MNIST and CIFAR10 dataset (each row depicts an example sequence), whereby a patch is shifted horizontally (left- or rightward) over the target image. **B**: Classification accuracy for models without (baseline) and with a temporal adaptation mechanism. **C-D**: Same as (B) for networks trained on fashion MNIST (C) and CIFAR10 (D) dataset. Recurrent adaptation mechanisms, both additive and multiplicative, show a gradual increase in performance. **E**: The top row shows the input sequence fed to a DCNN with multiplicative recurrent adaptation (*rec_A_*). The bottom row depicts the channel-averaged feature map of the first convolutional layer after adaptation is applied. Additive recurrent adaptation actively fills in occluded parts of the image with object-related features.

Initial comparison of performance across the sequence indicates that the baseline model and models with intrinsic adaptation exhibit a sudden drop in performance during the first few timesteps. These patterns were consistent across all three datasets, including MNIST (**Fig. 4B**), fashion MNIST (**Fig. 4C**) and CIFAR10 (**Fig. 4D**). The performance drop in the first few timesteps coincides with the patch covering more pixels belonging to the target, implying that the networks do not maintain a global reconstruction of the object over time as parts of it are occluded. In contrast, recurrent adaptation mechanisms show a gradual increase in performance as the patch slides over the object, suggesting that these mechanisms leverage the continuity of the patch’s motion and contextual information to “fill in” the occluded parts in the inputs. This is putatively due to the fact that the interactions across feature maps enable the network to facilitate a global reconstruction of the object over time, which leads to a smoother, more consistent improvement in performance compared to the baseline model and models with intrinsic adaptation. In support of this hypothesis, we observe for the recurrent adaptation mechanisms, in particular the additive form, a reconstruction of object-related features at the location of the patch after training on the MNIST and fashion MNIST dataset (**Fig. 4E**; for fashion MNIST and CIFAR10 see **Supp. Fig. 10**). This is possibly a result of combining information from different feature maps, enabling the network to reconstruct occluded parts in the image and refining its internal representation over time.

Notably, DCNNs with recurrent mechanisms show an initial lower performance at the beginning of the sequence in comparison with DCNNs endowed with intrinsic adaptation. Given that no prior temporal information is available at the first timestep for either type of mechanism, the difference must arise from the feedforward weights and how these were optimized during training in the presence of the different adaptation mechanisms. More specifically, these results strongly suggests an interaction effect between the optimization of feedforward parameters and the parameters modulating the temporal adaptation. We will return to the difference in initial performance between intrinsic and recurrent adaptation mechanisms in the Discussion. Nonetheless, these findings show that recurrent adaptation mechanisms excel at integrating spatio-temporal information over time leading to a smooth performance increase.

### Temporal adaptation mechanisms are required to detect novel objects

When a novel stimulus enters the visual field, attention is rapidly redirected towards it, a process known as novelty detection (Sokolov, 1963; Corbetta and Shulman, 2002). For instance, when driving on a road, a person’s focus might abruptly shift from the cars ahead to an unfamiliar road sign or an animal crossing the street. To investigate how the different temporal adaptation mechanisms contribute to the salience of novel visual inputs, we created image sequences whereby novel target images were presented over time and networks were trained to predict the object belonging to the most recently introduced target image (**Fig. 5A**; see Materials and Methods, *Experiment 3: Novelty detection*). Here, temporal adaptation could aid the detection of novel objects by allowing progressive adaptation to already familiar ones. Moreover, to assess whether adaptation mechanisms can differentiate between local changes in object appearance and global changes related to the presentation of novel objects, we applied a set of image augmentation techniques (e.g. rotation, scaling) to the target images at each model timestep. While DCNN instances were trained on all three datasets, the networks optimized on CIFAR10 failed to converge, and therefore, this dataset was excluded from the analysis.

**Figure 5:**
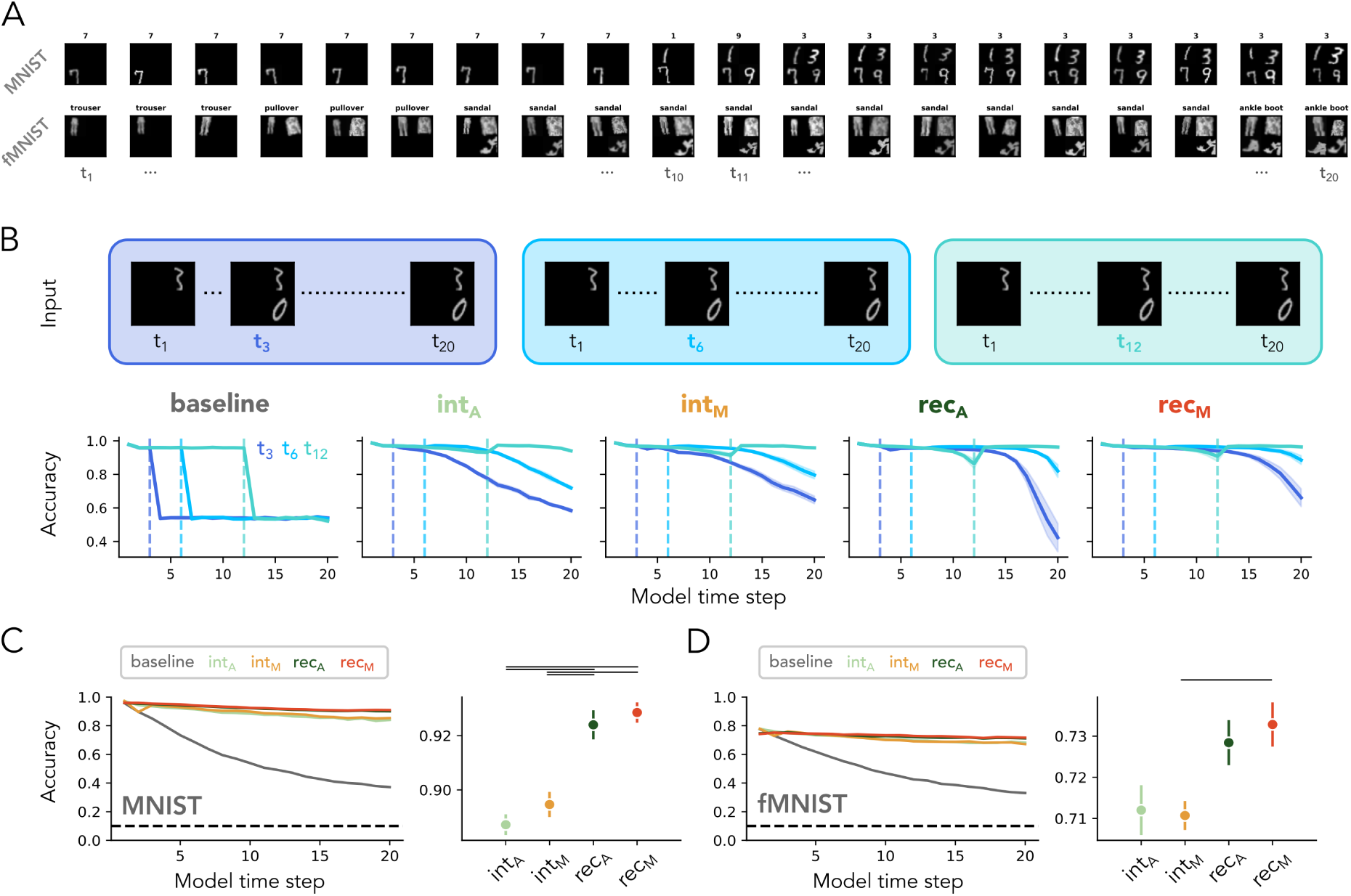
Recurrent adaptation mechanisms outperform intrinsic adaptation mechanisms during novelty detection. **A**: Each row depicts an example training sequence for the MNIST (top) and fashion MNIST (bottom) dataset with novel target images added over model timesteps. **B**: *Top*, Three input sequences fed to the network that contain two target images per sequence with the first target image presented on the first timestep and the second target image appearing on model timestep 3 (left), 6 (middle) or 12 (right). *Bottom*, Classification accuracy for models without (baseline) and with a temporal adaptation mechanism. Models with temporal adaptation are able to accurately classify the second object. **C**: The left panel shows the classification accuracy across model timesteps when trained on the MNIST dataset for a feedforward model without (baseline) and with a temporal adaptation mechanism. The right panel shows the classification accuracy averaged over model timesteps. Values depict means, error bars depict SEM across network initializations (*n* = 5) and the horizontal grey lines depict significant differences between networks (one-way ANOVA, post-hoc Tukey test, *p <* 0.05). **D**: Same as panel (B) but for networks trained on the fashion MNIST dataset. Recurrent adaptation mechanisms show higher performance compared to intrinsic adaptation mechanisms.

To investigate how the networks perform when novel objects are introduced, we first examined performance around the onset of a novel target image. For this, we created a simplified test set where instead of introducing four objects, as was done in the training set, we only present two target images, with the second target image introduced at timesteps 3, 6, or 12 (**Fig. 5B**). Baseline model performance drops instantly after the second target image is introduced, indicating that the model cannot know which of the two objects to classify. In contrast, accuracy of DCNNs with temporal adaptation remains high after presenting the second target image, demonstrating that the models use their temporal adaptation mechanism to enhance the salience of the novel object. We do notice a reduction in performance for later onsets (on *t*_3_ and *t*_6_), which is likely due to the fact that in the training sequences new target images were introduced with a relatively high rate (four target images in twenty model timesteps) and networks were therefore not optimized to classify the same target images for prolonged number of model timesteps, putatively leading to the observed decline in accuracy. Nonetheless, these findings show that temporal adaptation mechanisms aid in tracking novel objects among familiar ones.

Notably, we observe a qualitative difference across the different temporal adaptation mechanisms: performance of DCNNs with intrinsic adaptation deteriorates earlier in the sequence, while the accuracy of recurrent adaptation mechanisms remains high for a greater number of timesteps. For instance, for sequences with an onset of *t*_3_, accuracy starts to decline around timestep 6 for mechanisms with intrinsic adaptation, while for recurrent adaptation mechanisms, accuracy remains constant for at least 12 timesteps. Overall, these findings show that temporal adaptation allows DCNNs to detect novel objects in dynamic sequences and suggests that temporally integrating over feature maps, as is the case for recurrent adaptation mechanisms, aids in tracking novel objects more effectively.

### Recurrent adaptation exhibit higher performance during novelty detection

To examine whether the difference in performance between recurrent and intrinsic adaptation mechanisms is robust, we created sequences similar to those used in the training set, whereby a total of four target images were introduced and where the onset of each target image was varied. Networks without a temporal adaptation mechanism achieve an average accuracy of around 100% on the MNIST dataset and 80% on the fMNIST dataset when only one object is presented. However, performance rapidly declines as more images are added, confirming our previous observation that these networks struggle to detect novel objects. This pattern holds consistently across both datasets, namely MNIST (**Fig. 5C**) and fMNIST (**Fig. 5D**). We also confirm the observed differences across adaptation mechanisms, evident by the fact that DCNNs with recurrent adaptation outperform those with intrinsic adaptation (MNIST, one-way ANOVA, *F*_4_ = 1215.11, *p <* 0.001, **Supp. Table 6**; fMNIST, one-way ANOVA, *F*_4_ = 372.65, *p <* 0.001, **Supp. Table 7**). This improved performance is especially noticeable later in the sequence, when multiple target images are presented.

We hypothesize that that cross-feature map integration is beneficial for distinguishing between changes caused by the introduction of a new target image and those resulting from the augmentation of a familiar image that was presented earlier in the sequence. Intrinsic adaptation mechanisms which do not have this feature map integration, adjust their activations locally and therefore may have difficulty distinguishing between new target images and slight modifications of existing ones, especially when more target images are presented. To confirm this hypothesis, we also trained a set of DCNNs without applying augmentation to the target images, thereby removing the local variations over model timesteps. While recurrent adaptation still results in high DCNN performance, the performance difference between intrinsic adaptation, particularly the additive form, and recurrent adaptation disappears (**Supp. Fig. 11**), indeed suggesting that cross-feature map adaptation is particularly beneficial when the task requires to distinguish local changes from global changes related to novel incoming information. To conclude, these findings demonstrate that recurrent adaptation mechanisms show superior ability compared to intrinsic adaptation in tracking global changes allowing for the detection of novel objects.

## Discussion

The aim of this study was to understand how temporal adaptation may help achieve robust primate vision under suboptimal viewing scenario’s. By comparing four different temporal adaptation mechanisms implemented in a feedforward DCNN, we found that the impact of these mechanisms on object recognition depends on both the origin of the feedback signal (intrinsic vs. recurrent) and the type of interaction used to process previous inputs (additive vs. multiplicative), with their impact on performance interacting with the specific task and dataset. For object recognition under noise, intrinsic mechanisms are particularly effective at processing simple, high-contrast features, while recurrent mechanisms excel at handling complex, low-contrast features, with both mechanisms leading to an overall improvement in recognition performance compared to a baseline model without adaptation. During object recognition under occlusion, recurrent mechanisms excel at integrating spatio-temporal information over time, which leads to a smoother performance increase compared to intrinsic adaptation mechanisms. Finally, we show that during novelty detection, temporal adaptation mechanisms are crucial for detecting new objects, whereby recurrent adaptation mechanisms outperform intrinsic adaptation mechanisms by better distinguishing local from global changes related to novel inputs. Together, these results highlight different strengths of each type of temporal adaptation mechanism, suggesting that multiple mechanisms which are employed in parallel may be most effective for achieving robust object recognition in a variety of challenging real-world scenarios.

### Task design and dataset impact performance of temporal adaptation mechanisms

We find that temporal adaptation mechanisms can enhance a DCNN’s ability to recognize objects in a range of challenging scenario’s, including noise and occlusion, as well as in the context of novelty detection. Our study expands on prior research comparing feedforward DCNNs augmented with different computational mechanisms to gain insights into the role of temporal adaptation during object recognition (Vinken et al., 2020; Lindsay et al., 2022; Sörensen et al., 2023) by applying these comparisons to a broader range of tasks and datasets, while also introducing novel adaptation mechanisms (divisive normalization and multiplicative recurrence). This approach offers two valuable insights. First, we demonstrate that task design plays a crucial role in shaping model performance. For object recognition under noise, a model with additive intrinsic adaptation showed superior performance compared to the other adaptation mechanisms when noise was added at the last timestep only. However, the network did not generalize well to noise patterns that differed between the adapter and test images within a trial, suggesting that it learned to subtract the noise independently of the pattern. This contrasts with previous work in humans, which demonstrated that adapting to a different noise pattern can enhance performance compared to when no adapter is presented (Brands et al., 2024) and underscores the importance of designing tasks with sufficient complexity to ensure that specific components of the mechanism cannot be exploited in unintended ways. Second, by optimizing the networks on different tasks and datasets, we reveal that the performance benefit of a given temporal adaptation mechanism interacts with the specific design of the task and the features of the visual input being processed. This raises the interesting possibility that different temporal adaptation mechanisms serve distinct roles during object recognition and may be employed in parallel by the human brain, whereby the nature of object degradation, such as noise or occlusion, can significantly influence which mechanism is involved. Overall, these findings emphasize the necessity of careful consideration of task design and broad assessment across different viewing scenario’s and datasets to fully understand the role of different adaptation mechanisms on object recognition.

### Benefits of intrinsic and recurrent adaptation depends on input complexity

For object recognition under progressively decreasing noise, we show that DCNNs with intrinsic adaptation show higher performance when presented with simple, high-contrast inputs (MNIST) indicating that these mechanisms are sufficient to aid perception for these type of stimuli, as previously suggested by Vinken et al. (2020). Intrinsic adaptation mechanisms occurring on the level of the individual neuron have been observed across various brain regions and cell types in the visual cortex and have been implicated in early visual processing (e.g. Azouz and Gray 2000; Benda and Herz 2003; Mensi et al. 2012; for a review see Silver 2010; Whitmire and Stanley 2016). One possible implication of our findings is that intrinsic adaptation in early visual areas may play a role in quickly recognizing high-contrast features, such as edges and orientation, which are relatively easy to distinguish from temporally repeated noise and are indeed sufficient for robust object perception.

When presented with more complex inputs (fashion MNIST and CIFAR10), networks benefited most from interactions across feature maps, evident by the fact that recurrent adaptation resulted in higher DCNN performance compared to intrinsic adaptation, which implies that interactions within the same layer benefit object recognition when there is more variance in contrasts and textures. In biological systems, there is evidence of adaptation across circuits, for example during stimulus-specific adaptation (Ulanovsky et al., 2003; Sawamura et al., 2006) and adaptation of orientation tuning (Dragoi et al., 2000; Felsen et al., 2002) and this has also been shown to influence perception (Anstis et al., 1998; Zavitz et al., 2016). Our results imply that these types of adaptive processes within neural circuits are required for object recognition when object features are less pronounced and differentiation between noise and object is more challenging.

Overall, our results suggest that intrinsic mechanisms operating on the individual neuron level and recurrent processes operating across neural circuits may have distinct roles when detecting objects from noise in a dynamic environment depending on the complexity of the visual input.

### Additive modulation benefits object recognition during dynamic occlusion

Our occlusion experiments reveal differences in how computational mechanisms integrate the visual inputs over time and how they process obstructed parts in the image. First, we find that recurrent adaptation mechanisms exhibit a smooth increase in performance as a patch slides across the target image. This progressive increase implies that the integration across feature maps aids in constructing a coherent representation of the partially visible object and is supported by the fact that recurrent adaptation leads to the reconstruction of object-related features in the occluded parts of the image. The nature of object representations in biological systems during occlusion has been examined using a variety of approaches, where some studies have suggested the representation during occlusion is an abstract representation of the object’s position only (Pylyshyn, 2004; Flombaum et al., 2004). In contrast, other studies have suggested there may be a richer object representation present, which contains information about object shape, category and identity (Hulme and Zeki, 2007; Puneeth et al., 2016). These contrasting views highlight the complexity of the nature of the object representation during occlusion, whereby our results suggest that recurrent adaptation mechanisms might be more aligned with richer, dynamic representations of objects, rather than abstract ones.

We also observe an initial performance difference across adaptation mechanisms, with intrinsic adaptation mechanisms resulting in a higher network performance at the first timestep compared to recurrent mechanisms. Since no prior temporal information is available at the first timestep, the difference must arise from the feedforward weights. Moreover, the fact that the feedforward weights were optimized together with the temporal adaptation parameters, suggests that there is an interaction effect between the different types of parameters, which influences the convergence during training. Previous work has attempted to tackle this and isolate the effect of different temporal adaptation mechanisms by using a pre-trained feedforward DCNN and solely optimizing the parameters of the adaptation mechanisms (Lindsay et al., 2022). Follow-up research could deploy a similar approach, allowing to directly compare the impact of different types of temporal adaptation mechanisms and how they aid object recognition in motion occluding paradigms.

### Recurrent adaptation tracks global changes during novelty detection

We have shown that temporal adaptation mechanisms aid in detecting novel objects by adapting, and thereby reducing the salience of familiar ones. This finding is in line with previous work in visual cortex, which has shown that distinguishing novel and familiar images is a result of adaptation processes resulting in increased responses to novel visual stimuli (Vinken et al., 2017; Homann et al., 2022). We also show that recurrent adaptation mechanisms are more effective at distinguishing between local changes caused by small alterations in object appearance and global changes caused by the presentation of novel information, leading to better recognition performance compared to intrinsic adaptation mechanisms. In early visual cortex, individual neurons process and adapt to local patches in the image, determined by their receptive field (Hubel and Wiesel, 1962), and our findings suggest that this local adaptation allows for the detection of novel objects. However, our results also imply that more fine-grained differentiation between local and global changes entails context-dependent modulation as a result of recurrent connections within larger neural circuits. Previous work has related lateral recurrent to certain aspects of visual processing, such as refining feature extraction (Neumann and Sepp, 1999; Niell and Scanziani, 2021) and encoding sensory variance (Ladret et al., 2023). On the perceptual level, earlier research has also showed that recurrent processing in early visual areas correlates with attention and awareness to visual inputs (Boehler et al., 2008). These findings suggest that recurrent processing plays a key role in enhancing perceptual sensitivity by refining responses to both local and global features of the visual scene.

### Limitations and future work

First, future comparisons to biological systems in the form of neural single- or multi-cell recordings, in combination with the collection of behavior, will be necessary to investigate the role of different temporal adaptation mechanisms on perception. Second, we intentionally used a more lightweight DCNN architecture in this study, with temporal adaptation being applied only in the first convolutional network layer, to make analysis of network behavior and activity more tractable. However, such an approach has limitations when comparing it to complex biological systems, which contains substantially more units and likely implements adaptation across multiple hierarchical levels. Training larger network architectures and implementing adaptation throughout the network are therefore important future directions for teasing apart the role of temporal adaptation in the human visual system. Third, previous work on understanding classification has pointed to the role of auxiliary variables, referring to image features not directly relevant to the classification such as object size and pose, as relevant pieces of information for the network to keep track of (Hong et al., 2016; Thorat et al., 2021). Future research could further investigate how different mechanisms represent such fea-tures over over time and the implications on robust object recognition. Fourth and last, while in the current study we implemented just one of four computational mechanisms in a single DCNN at a time, our results imply that biological system perform a combination of these mechanisms simultaneously. A fruitful endeavor would therefore be to study how a combination of both intrinsic and recurrent adaptation mechanisms jointly affect object recognition and aid in robust perception under challenging viewing scenarios.

---

**Supplementary Figure 1:**
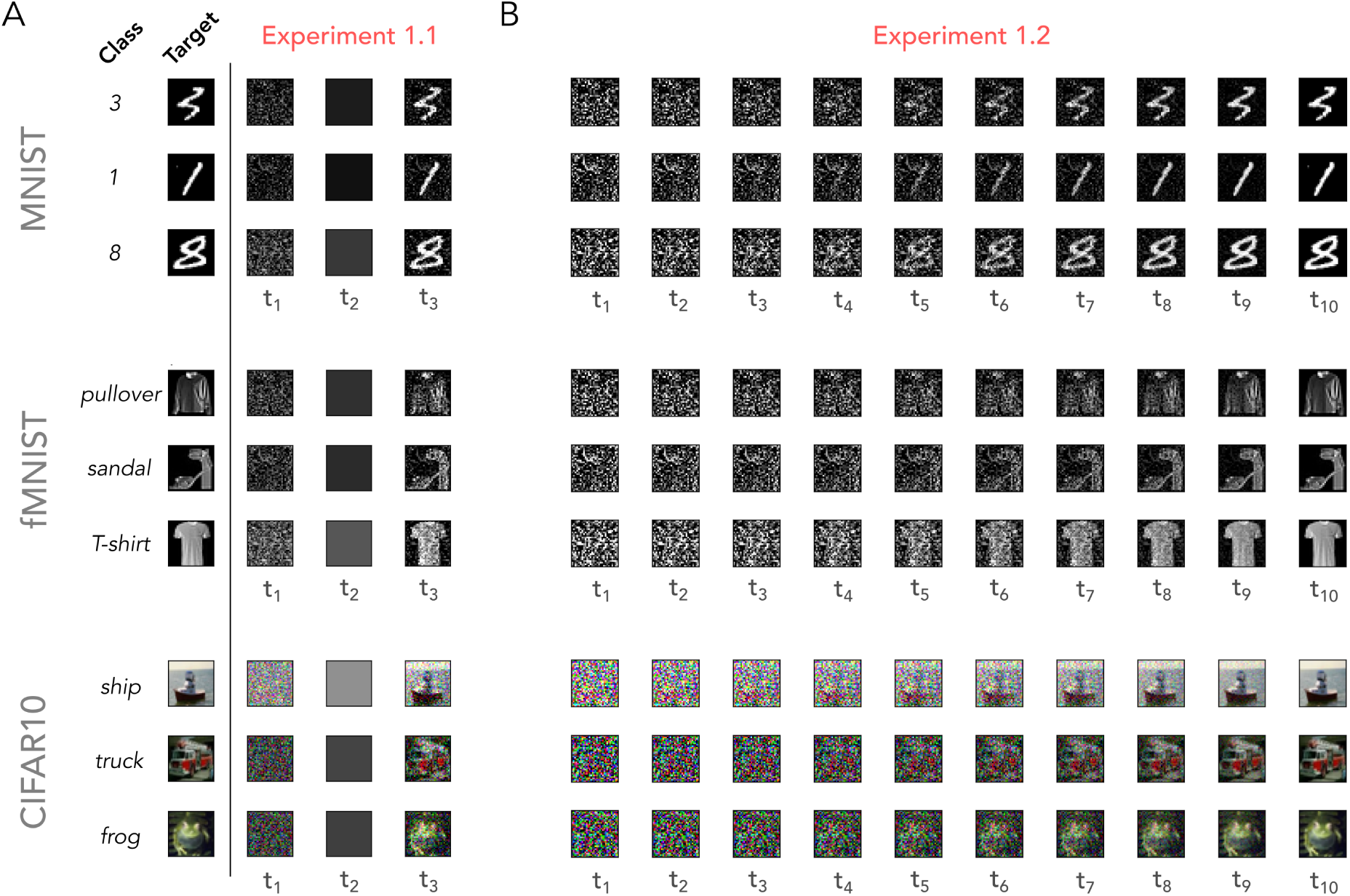
Overview of Experiment 1 (Object recognition under noise). **A**: *Experiment 1.1*, Input sequences contained a target image with an object belonging to one of ten classes from the MNIST, fashion MNIST (fMNIST) or CIFAR10 dataset. Input sequences (*t* = 3) consisted of three model timesteps. First, an adapter image was shown (*t*_1_), followed by a blank image (*t*_2_) and a test image (*t*_3_), which consisted of the target image embedded in the same noise pattern as was used for the adapter. For each dataset, three example sequences are shown (rows). **B**: *Experiment 1.2*, Input sequences (*t* = 10) consisted of a target image superimposed on Gaussian noise, whereby the ratio between the target image and noise was increased with every model time step. For each dataset, three example sequences are shown (rows).

**Supplementary Figure 2:**
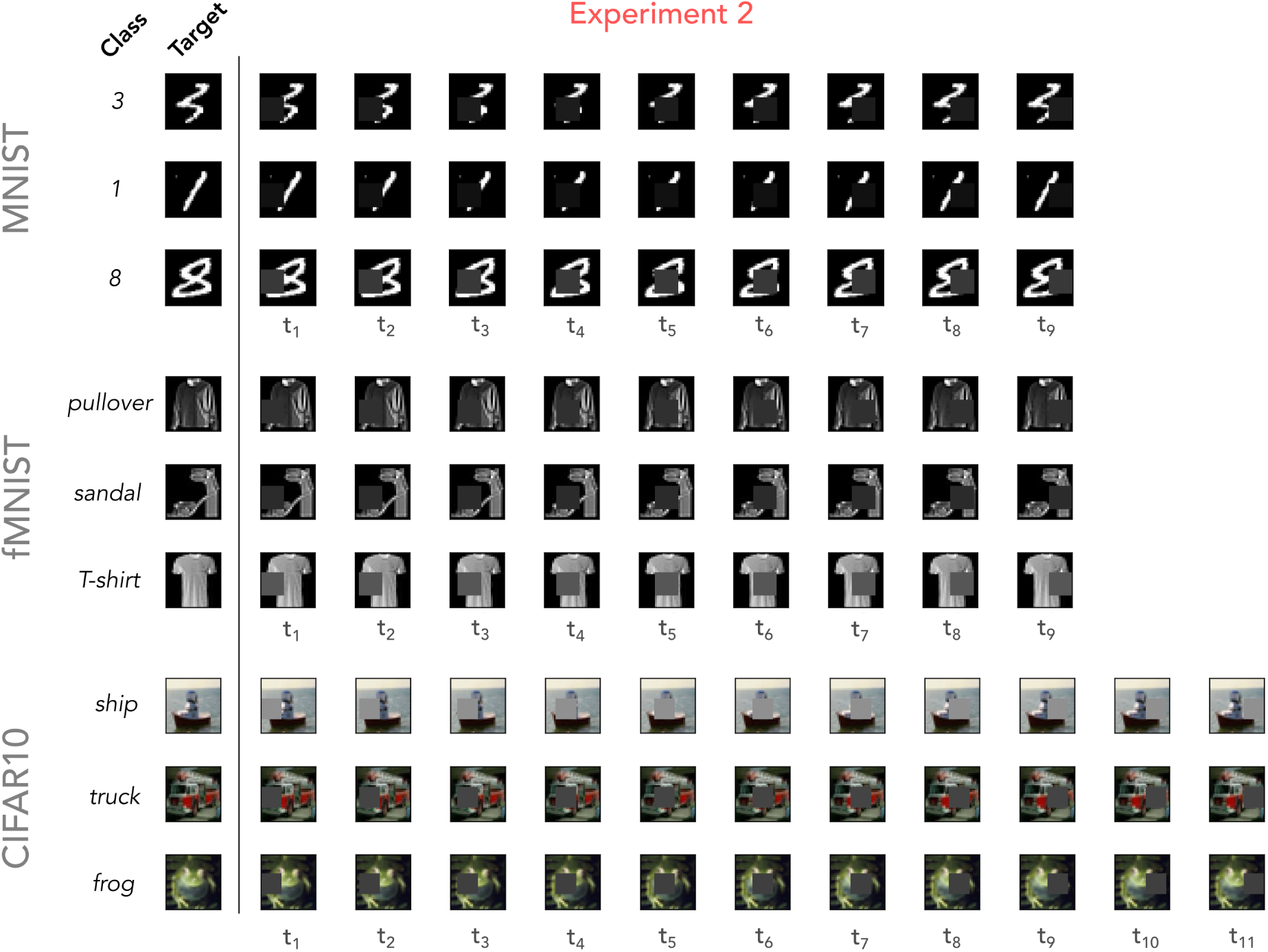
Overview of Experiment 2 (Object recognition under occlusion). Input sequences contained a target image with an object belonging to one of ten classes of the MNIST (*t* = 9), fashion MNIST (fMNIST, *t* = 9) or CIFAR10 (*t* = 11) dataset. With every model timestep, a patch was shifted two pixels to the left or right (depending on whether the starting position of the patch was right or left, resp.), thereby occluding the object. For each dataset, three example sequences are shown (rows).

**Supplementary Figure 3:**
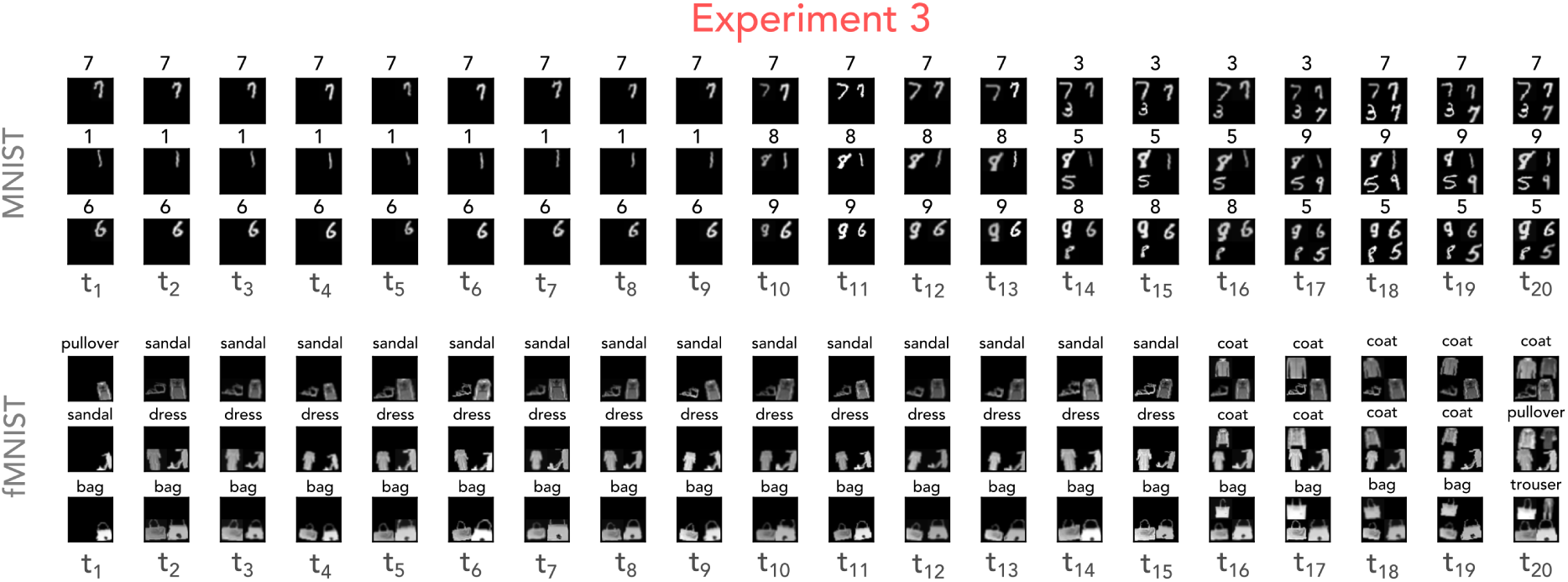
Overview of Experiment (Novelty detection). Input sequences (*t* = 20) consisted of an image, whereby target images with an object belonging to one of ten classes of the MNIST or fashion MNIST (fMNIST) dataset were randomly added over model time steps. For each timestep and object, augmentation techniques were applied (e.g. rotation, scaling, translation, varying contrast). For each dataset, three example sequences are shown (rows). Above each image in the sequence, the ground truth label is reported.

**Supplementary Figure 4:**
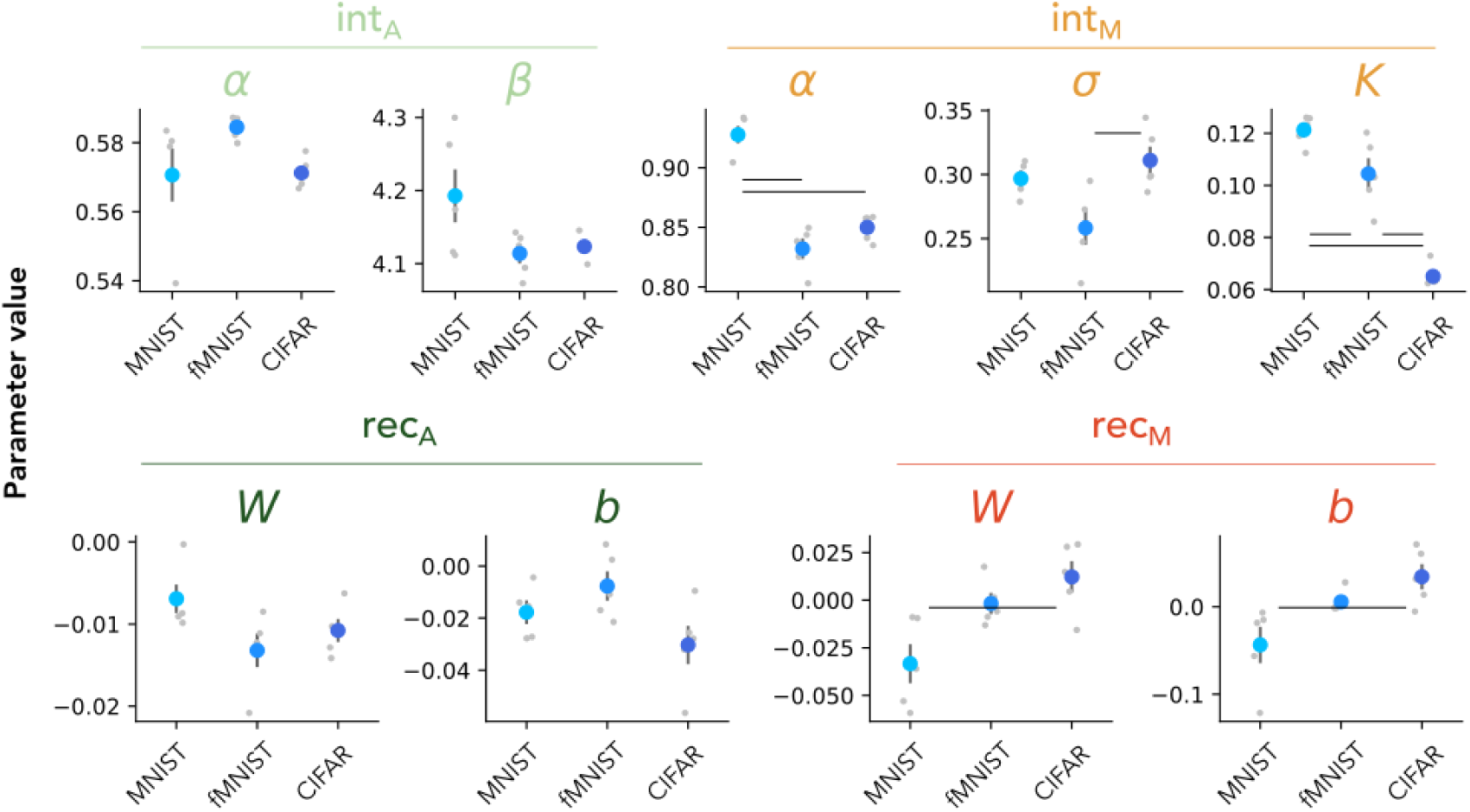
Fitted parameters for Experiment 1.1 (Object recognition under noise). Trained parameter values for a feedforward model with a temporal adaptation mechanism, arising from intrinsic (int) or recurrent (rec) mechanisms, using additive (A) or multiplicative (M) interactions, averaged over network initializations (*n* = 5) per dataset, including MNIST, fashion MNIST (fMNIST) and CIFAR10 (CIFAR). Individual points depict values for individual network initializations and error bars depict standard deviations across initializations.

**Supplementary Figure 5:**
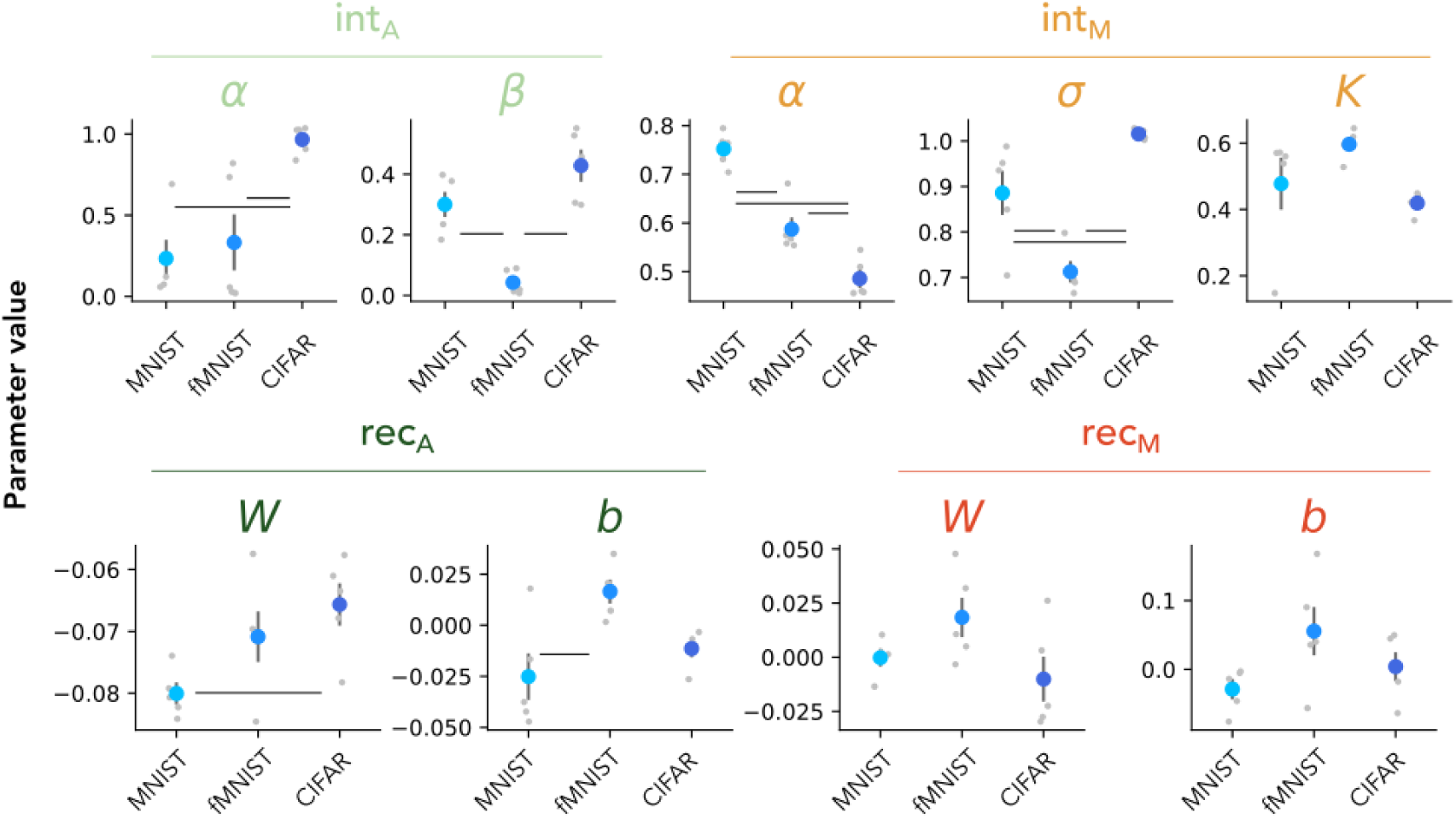
Fitted parameters for Experiment 1.2 (Object recognition under noise). Trained parameter values for a feedforward model with a temporal adaptation mechanism, arising from intrinsic (int) or recurrent (rec) mechanisms, using additive (A) or multiplicative (M) interactions, averaged over network initializations (*n* = 5) per dataset, including MNIST, fashion MNIST (fMNIST) and CIFAR10 (CIFAR). Individual points depict values for individual network initializations and error bars depict standard deviations across initializations.

**Supplementary Figure 6:**
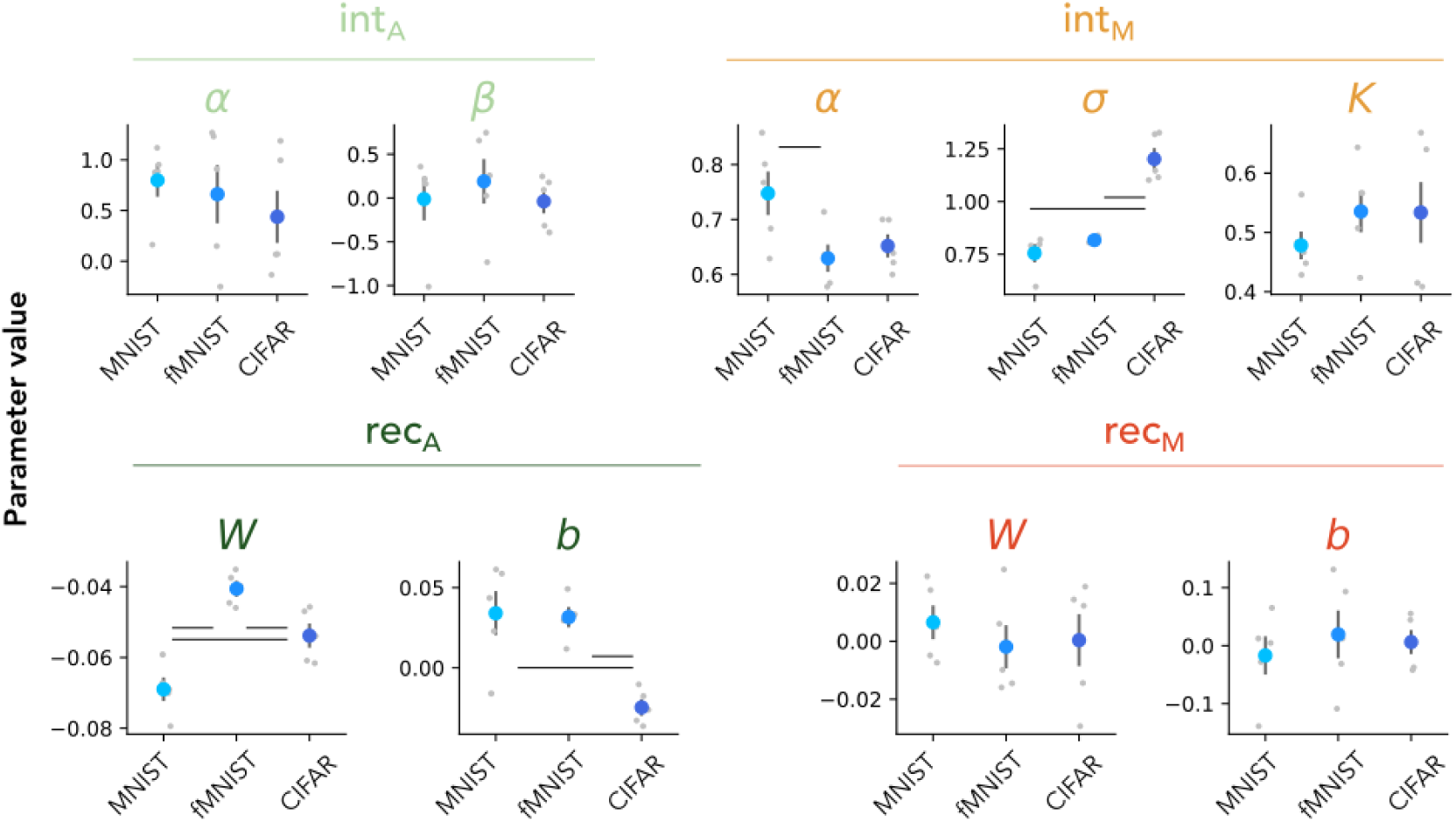
Fitted parameters for Experiment 2 (Object recognition under occlusion). Trained parameter values for a feedforward model with a temporal adaptation mechanism, arising from intrinsic (int) or recurrent (rec) mechanisms, using additive (A) or multiplicative (M) interactions, averaged over network initializations (*n* = 5) per dataset, including MNIST, fashion MNIST (fMNIST) and CIFAR10 (CIFAR). Individual points depict values for individual network initializations and error bars depict standard deviations across initializations.

**Supplementary Figure 7:**
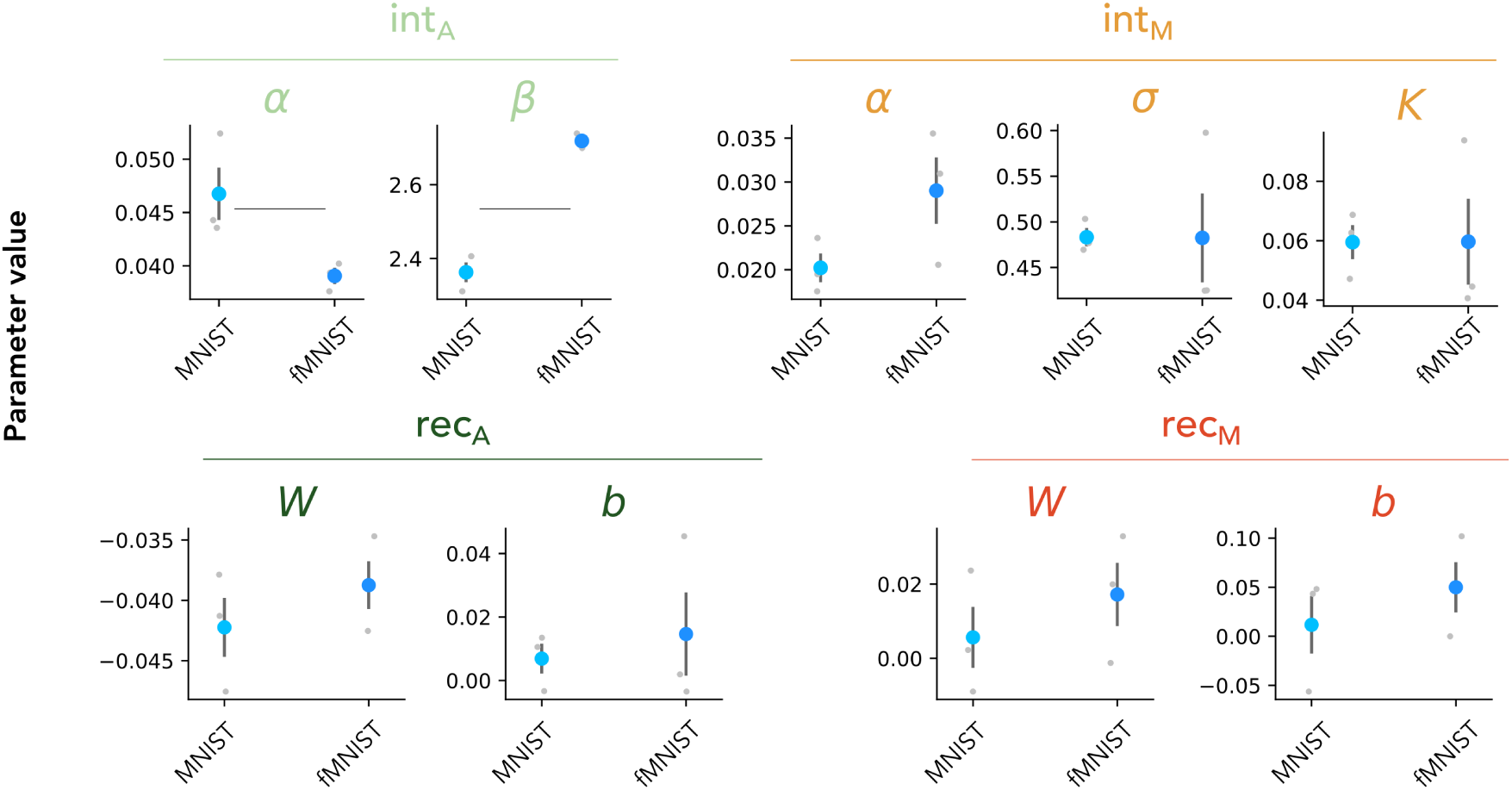
Fitted parameters for Experiment 3 (Novelty detection). Trained parameter values for a feedforward model with a temporal adaptation mechanism, arising from intrinsic (int) or recurrent (rec) mechanisms, using additive (A) or multiplicative (M) interactions, averaged over network initializations (*n* = 5) per dataset, including MNIST, fashion MNIST (fMNIST) and CIFAR10 (CIFAR). Individual points depict values for individual network initializations and error bars depict standard deviations across initializations.

**Supplementary Figure 8:**
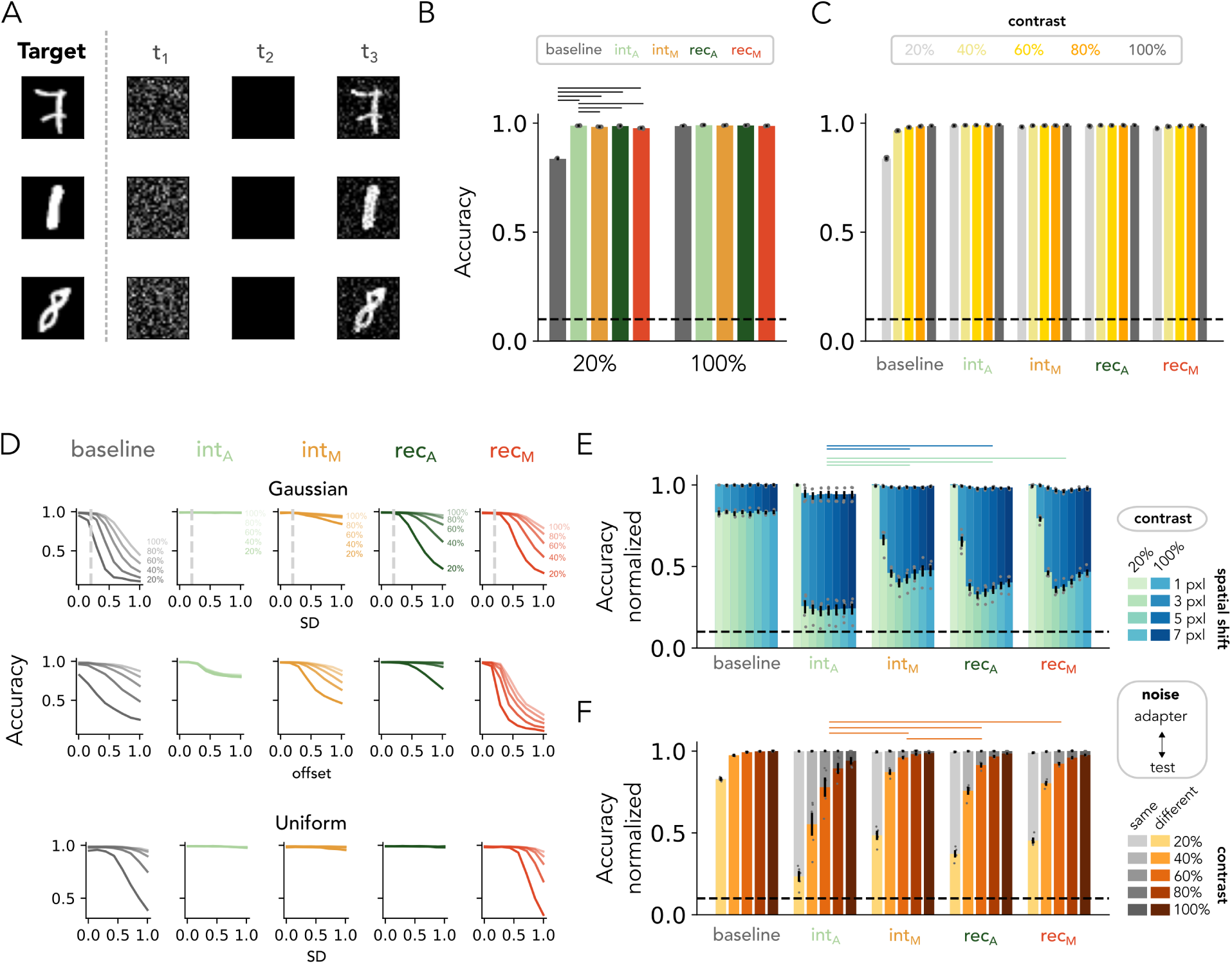
DCNNs with additive intrinsic adaptation solves task by subtracting the noise. **A**: Each row depicts an example sequence. The first image in the sequence (*t*_1_) contains Gaussian noise and is followed by a blank image (*t*_2_). The third image (*t*_3_) contains a MNIST target image embedded in the noise pattern presented at (*t*_1_). **B**: Test accuracy for models with and without (*baseline*) temporal adaptation were trained on a mix of high- (100 %) and low- (20%) object contrasts. DCNNs with additive intrinsic adaptation (*int_A_*) exhibit highest performance for both object contrast levels. **C**: Model performance when generalizing to different object contrasts (only the contrast levels in gray were included in the training set). Superior performance of DCNNs with additive intrinsic adaptation (*int_A_*) consistent across object contrast levels. **D**: Model performance when generalizing to different noise patterns (see Materials and Methods, *Experiment 1: Object recognition under noise*). Superior performance of DCNNs with additive intrinsic adaptation (*int_A_*) consistent across different types of noise. **E**: Model performance when spatially shifting the noise pattern during test. The first bar for each adaptation mechanism depicts performance without a spatial shift. **F**: Model performance when the test image (*t*_3_) contains a different noise pattern than the adapter image (*t*_1_). Grey bars mark accuracy when tested on the same noise pattern (same data as shown in panel C). Superior performance of DCNNs with additive intrinsic adaptation (*int_A_*) achieved by subtraction of the noise, resulting in lower robustness to spatial shifts (E) and different noise during the test (F). For panel B, C, E and F, the dashed line depicts chance level (i.e. 10%), markers represent individual network initializations and the error bars depict the SEM across these network initializations. For panel B, E and F, horizontal stripes on the top of the plot indicate significant differences between networks (One-way ANOVA, post-hoc Tukey test, *p <* 0.05).

**Supplementary Figure 9:**
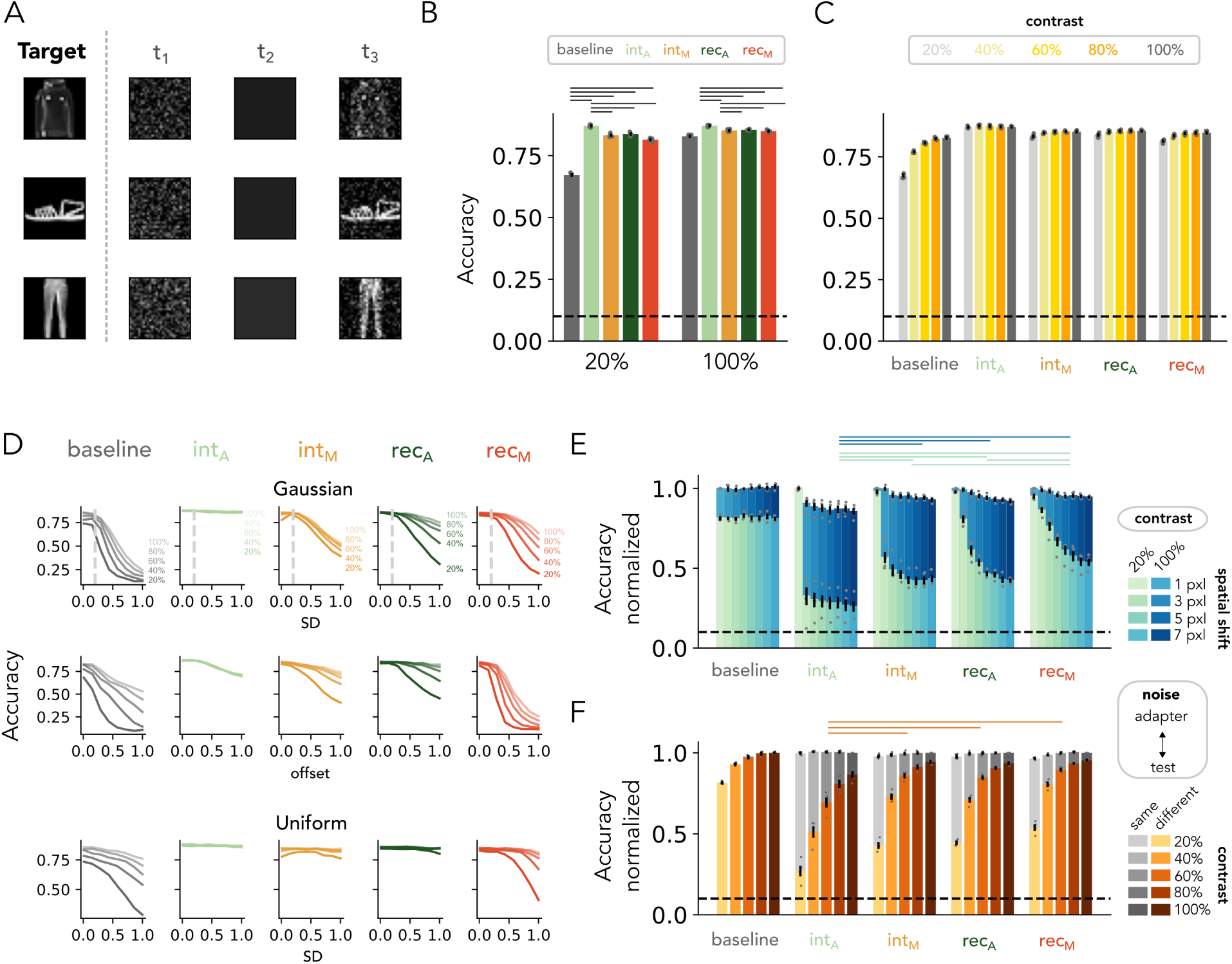
DCNNs with additive intrinsic adaptation solves task by subtracting the noise. **A**: Each row depicts an example sequence. The first image in the sequence (*t*_1_) contains Gaussian noise and is followed by a blank image (*t*_2_). The third image (*t*_3_) contains a fashion MNIST target image embedded in the noise pattern presented at (*t*_1_). **B**: Test accuracy for models with and without (*baseline*) temporal adaptation were trained on a mix of high- (100 %) and low- (20%) object contrasts. DCNNs with additive intrinsic adaptation (*int_A_*) exhibit highest performance for both object contrast levels. **C**: Model performance when generalizing to different object contrasts (only the contrast levels in gray were included in the training set). Superior performance of DCNNs with additive intrinsic adaptation (*int_A_*) consistent across object contrast levels. **D**: Model performance when generalizing to different noise patterns (see Materials and Methods, *Experiment 1: Object recognition under noise*). Superior performance of DCNNs with additive intrinsic adaptation (*int_A_*) consistent across different types of noise. **E**: Model performance when spatially shifting the noise pattern during test. The first bar for each adaptation mechanism depicts performance without a spatial shift. **F**: Model performance when the test image (*t*_3_) contains a different noise pattern than the adapter image (*t*_1_). Grey bars mark accuracy when tested on the same noise pattern (same data as shown in panel C). Superior performance of DCNNs with additive intrinsic adaptation (*int_A_*) achieved by subtraction of the noise, resulting in lower robustness to spatial shifts (E) and different noise during the test (F). For panel B, C, E and F, the dashed line depicts chance level (i.e. 10%), markers represent individual network initializations and the error bars depict the SEM across these network initializations. For panel B, E and F, horizontal stripes on the top of the plot indicate significant differences between networks (One-way ANOVA, post-hoc Tukey test, *p <* 0.05).

**Supplementary Figure 10:**
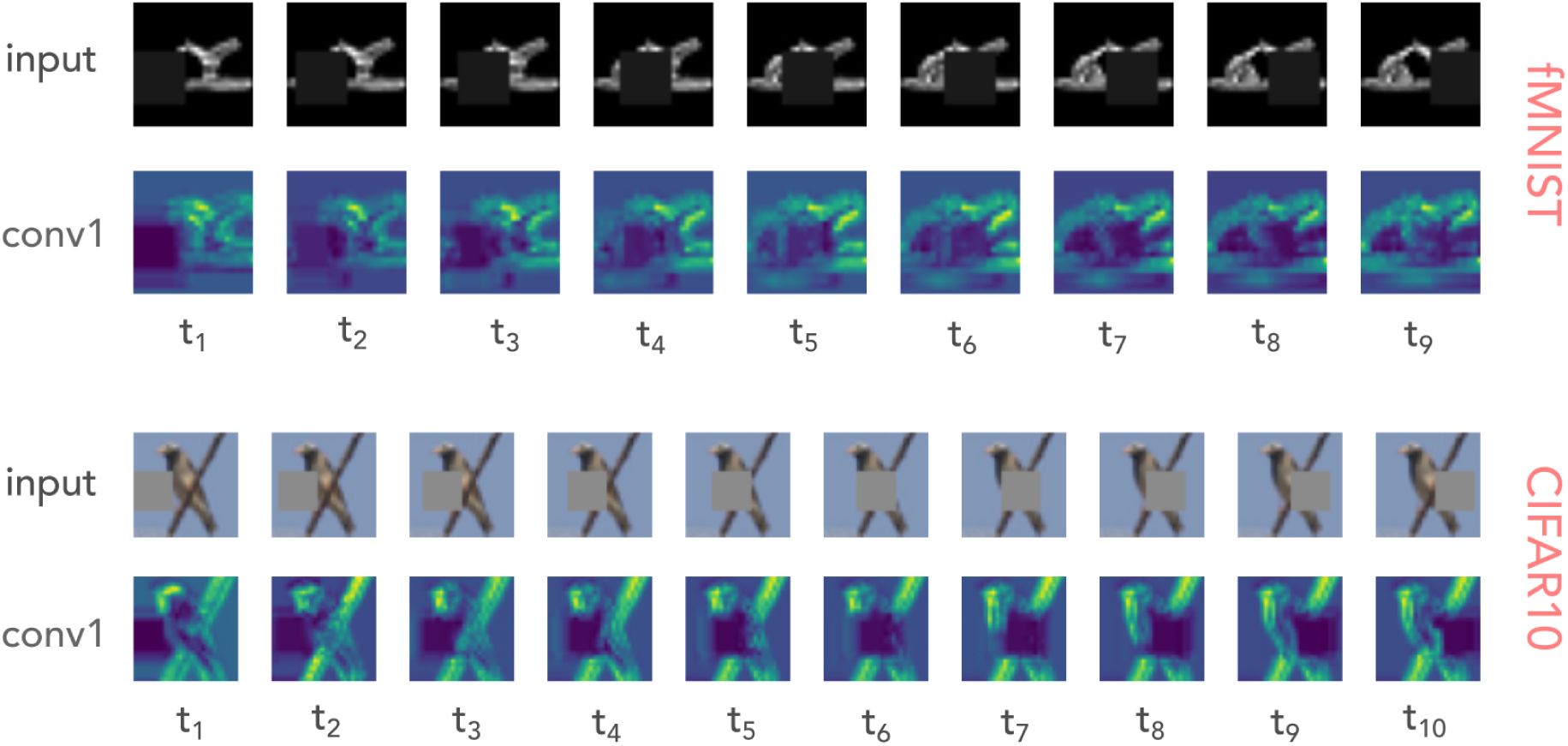
Feature maps for the first convolutional layer during object recognition under occlusion. Sample sequences for target images belonging the fashion MNIST (top) or CIFAR10 (bottom) dataset. For each dataset, the top row shows the input sequence fed to a DCNN with multiplicative recurrent adaptation (*rec_A_*). The bottom row depicts the channel-averaged feature map of the first convolutional layer after adaptation is applied.

**Supplementary Figure 11:**
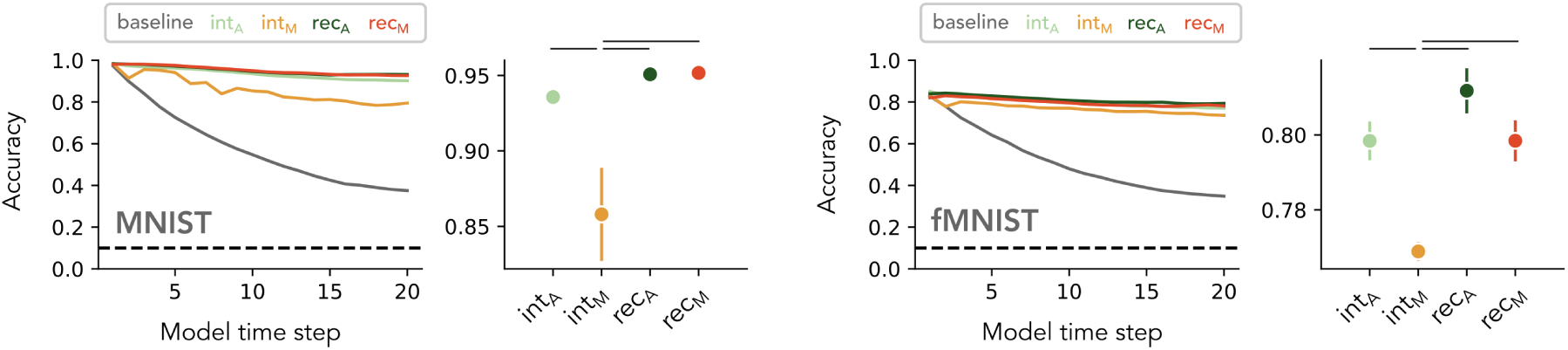
Performance during novelty detection without image augmentation. Performances for networks trained on target images belonging the fashion MNIST (left) or fashion MNIST (right) dataset. For each dataset, the left panel shows the classification accuracy across model timesteps for a feedforward model without (baseline) and with a temporal adaptation mechanism. The right panel shows the classification accuracy averaged over model timesteps. Values depict means, error bars depict SEM across network initializations (*n* = 5) and the horizontal gray lines depict significant differences between networks (one-way ANOVA, post-hoc Tukey test, *p <* 0.05).

**Supplementary Table 1:** Details of image augmentations applied to the target images.

| Augmentation | Description |
| --- | --- |
| Rotation | Objects were subjected to rotations, varying between -12 and 12 degrees. |
| Translation | A random uniform translation was applied in both x and y directions, ranging from 0 to 4 pixels. |
| Scaling | Objects underwent random uniform scaling, with a multiplication factor between 0.9 and 1.1. |
| Shear | Random shear transformations were applied, with angles from -5 to 5 degrees. |
| Gaussian noise | Gaussian noise, with a standard deviation of 0.1 (in pixel intensity values), was added to the images. |
| Brightness adjustment | The brightness of objects was altered, with adjustment factors from 0.8 to 1.2 (multiplier applied to pixel intensity values). |
| Contrast adjustment | Contrast was randomly adjusted, with factors ranging from 0.8 to 1.2 (multiplier applied to the pixel contrast). |

**Supplementary Table 2:** Model architectures for each experiment and dataset. Each row in each table represents a layer. *H*, *W* and *C* specifies the input size, namely height, width and number of channels, respectively. *F* specifies the number of feature maps in the layer, *K* represents the height and width of the convolutional kernel and *S* the stride used for the max pooling operation. All convolutions are applied with 1 × 1 stride. Dropout was set to 50%. The values in the linear layers depict the output size, where the output size of the last layer equals the number of classes in the datasets (i.e. 10).

| Experiment 1 (noise) and Experiment 2 (occlusion) |  |  |  | Experiment 3 (novelty detection) |  |  |  |
| --- | --- | --- | --- | --- | --- | --- | --- |
|  | MNIST | fMNIST | CIFAR-10 |  | MNIST | fMNIST | CIFAR-10 |
| *input* | H, W = 28, C = 1 | H, W = 28, C = 1 | H, W = 32, C = 3 | *input* | H, W = 56, C = 1 | H, W = 56, C = 1 | H, W = 64, C = 3 |
| conv2d | F = 32, K = 5 | F = 32, K = 5 | F = 32, K = 5 | conv2d | F = 32, K = 5 | F = 32, K = 5 | F = 32, K = 3 |
| ReLU | - | - | - | ReLU | - | - | - |
| MaxPool2d | K = 2, S = 2 | K = 2, S = 2 | K = 2, S = 2 | MaxPool2d | K = 2, S = 2 | K = 2, S = 2 | K = 2, S = 2 |
| conv2d | F = 32, K = 5 | F = 32, K = 5 | F = 64, K = 5 | conv2d | F = 32, K = 5 | F = 32, K = 5 | F = 64, K = 3 |
| ReLU | - | - | - | ReLU | - | - | - |
| MaxPool2d | K = 2, S = 2 | K = 2, S = 2 | K = 2, S = 2 | MaxPool2d | K = 2, S = 2 | K = 2, S = 2 | K = 2, S = 2 |
| conv2d | F = 32, K = 3 | F = 32, K = 3 | F = 64, K = 3 | conv2d | F = 32, K = 5 | F = 32, K = 5 | F = 64, K = 3 |
| ReLU | - | - | - | ReLU | - | - | - |
| Dropout | - | - | - | conv2d | F = 32, K = 3 | F = 32, K = 3 | F = 32, K = 3 |
| Linear | 1024 | 1024 | 1024 | ReLU | - | - | - |
| Linear | 10 | 10 | 10 | Dropout | - | - | - |
|  |  |  |  | Linear | 1024 | 1024 | 1024 |
|  |  |  |  | Linear | 10 | 10 | 10 |

**Supplementary Table 3:**
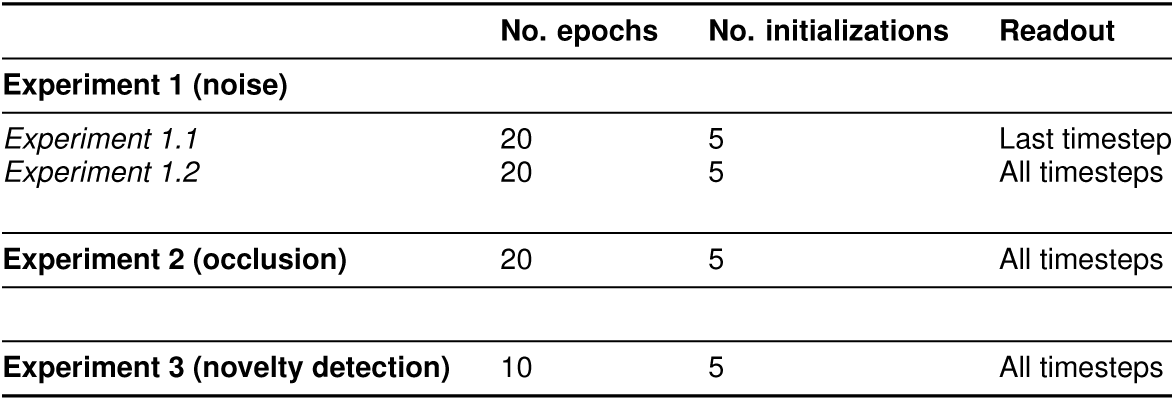
Training procedures per experiment. Columns refer to the following: *No. epochs*, The number of epochs networks were trained on. *No. initializations*, Number of networks that were trained with randomly initialized weights and biases. *Readout*, Time step which was used for class prediction.

**Supplementary Table 4:**
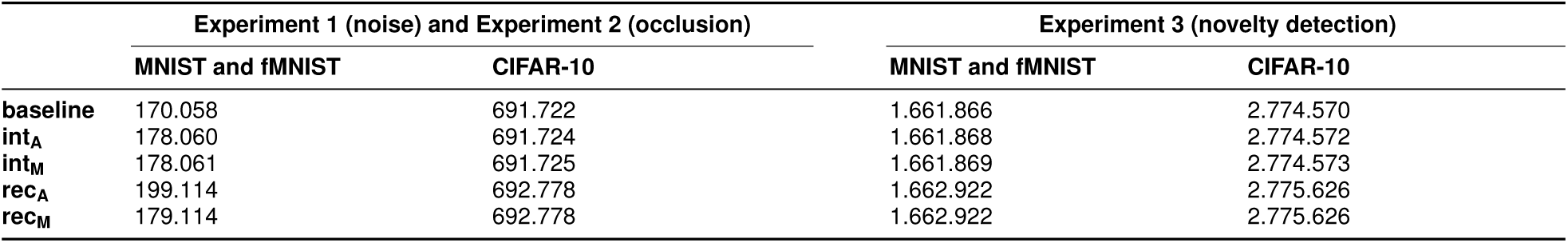
Number of trainable parameters per model, experiment and dataset. Shown are the number of parameters for a feedforward model without (none) and with a temporal adaptation, arising from intrinsic (int) or recurrent (rec) mechanisms, using additive (A) or multiplicative (M) interactions separately for each dataset.

**Supplementary Table 5:** Model evaluation on CIFAR-10 dataset. Classification accuracy (mean±SD) on the test set for a feedforward model without (baseline) and with temporal adaptation, arising from intrinsic (int) or recurrent (rec) mechanisms, using additive (A) or multiplicative (M) interactions.

|  | CIFAR-10 |  |  |  |  |
| --- | --- | --- | --- | --- | --- |
|  | baseline | int<sub>A</sub> | int<sub>M</sub> | rec<sub>A</sub> | rec<sub>M</sub> |
| **Experiment 1 (noise)** |  |  |  |  |  |
| Experiment 1.1 - *Low contrast* | 34.01±1.07 | 63.44±0.89 | 49.98±1.87 | 51.54±3.33 | 46.16±1.21 |
| Experiment 1.1 - *High contrast* | 52.74±1.50 | 63.13±1.04 | 55.53±0.95 | 56.34±1.79 | 55.30±0.51 |
| *Experiment 1.2* | 18.87±1.36 | 19.67±1.14 | 19.65±1.14 | 19.04±1.22 | 20.91±1.41 |
| **Experiment 2 (occlusion)** |  |  |  |  |  |
| *Experiment 2* | 53.52±1.83 | 53.29±0.88 | 42.35±12.44 | 50.61±1.36 | 42.45±2.38 |

**Supplementary Table 6:** Model evaluation on MNIST dataset. Classification accuracy on the test set for a feedforward model without (baseline) and with temporal adaptation, arising from intrinsic (int) or recurrent (rec) mechanisms, using additive (A) or multiplicative (M) interactions.

|  | MNIST |  |  |  |  |
| --- | --- | --- | --- | --- | --- |
|  | baseline | int<sub>A</sub> | int<sub>M</sub> | rec<sub>A</sub> | rec<sub>M</sub> |
| **Experiment 1 (noise)** |  |  |  |  |  |
| Experiment 1.1 - *Low contrast* | 81.59±0.86 | 99.01±0.09 | 98.32±0.15 | 98.59±0.19 | 97.63±0.18 |
| Experiment 1.1 - *High contrast* | 98.49±0.16 | 99.08±0.07 | 98.82±0.10 | 98.94±0.12 | 98.64±0.14 |
| *Experiment 1.2* | 56.93±2.37 | 58.82±1.78 | 58.33±2.34 | 55.47±2.88 | 53.48±4.75 |
| **Experiment 2 (occlusion)** |  |  |  |  |  |
| *Experiment 2* | 78.82±1.57 | 78.77±5.01 | 70.09±2.11 | 80.45±3.53 | 66.86±2.84 |
| **Experiment 3 (novelty detection)** |  |  |  |  |  |
| *Experiment 3* | 57.42±1.21 | 93.66±0.62 | 87.04±6.22 | 95.42±0.34 | 95.37±0.45 |

**Supplementary Table 7:** Model evaluation on fashion MNIST dataset. Classification accuracy (mean±SD) on the test set for a feedforward model without (baseline) and with temporal adaptation, arising from intrinsic (int) or recurrent (rec) mechanisms, using additive (A) or multiplicative (M) interactions.

|  | fashion MNIST |  |  |  |  |
| --- | --- | --- | --- | --- | --- |
|  | baseline | int<sub>A</sub> | int<sub>M</sub> | rec<sub>A</sub> | rec<sub>M</sub> |
| **Experiment 1 (noise)** |  |  |  |  |  |
| Experiment 1.1 - *High contrast* | 67.17±0.69 | 87.04±0.49 | 83.25±0.61 | 83.79±0.68 | 81.51±0.55 |
| Experiment 1.1 - *Low contrast* | 82.85±0.58 | 87.09±0.37 | 85.21±0.48 | 85.43±0.33 | 84.91±0.55 |
| *Experiment 1.2* | 43.02±1.46 | 42.63±1.29 | 43.88±1.26 | 45.49±1.27 | 44.52±1.21 |
| **Experiment 2 (occlusion)** |  |  |  |  |  |
| *Experiment 2* | 72.34±1.92 | 71.69±9.41 | 67.62±1.94 | 70.13±3.17 | 61.80±2.61 |
| **Experiment 3 (novelty detection)** |  |  |  |  |  |
| *Experiment 3* | 49.57±0.84 | 71.20±1.29 | 71.07±0.72 | 72.84±1.16 | 73.28±1.14 |

